# Behavioral Consequences of Velocity Commands: Brownian Processes in Human Motor Tasks

**DOI:** 10.1101/2023.05.17.541137

**Authors:** Federico Tessari, James Hermus, Rika Sugimoto-Dimitrova, Neville Hogan

**Affiliations:** Department of Mechanical Engineering, Massachusetts Institute of Technology, Cambridge, Massachusetts, USA; Department of Brain and Cognitive Sciences, Massachusetts Institute of Technology, Cambridge, Massachusetts, USA

## Abstract

The motor neuroscience literature suggests that the central nervous system may encode some motor commands in terms of velocity. In this work, we tackle the question: what consequences would velocity commands produce at the behavioral level? Considering the ubiquitous presence of noise in the neuromusculoskeletal system, we predict that velocity commands affected by stationary noise would produce “random walks”, also known as Brownian processes, in position. Brownian motions are distinctively characterized by a linearly growing variance and a power spectral density that declines in inverse proportion to frequency. This work first shows that these Brownian processes are indeed observed in unbounded motion tasks e.g., rotating a crank. We further predict that such growing variance would still be present, but bounded, in tasks requiring a constant posture e.g., maintaining a static hand position or quietly standing. This hypothesis was also confirmed by experimental observations. A series of descriptive models are investigated to justify the observed behavior. Interestingly, one of the models capable of accounting for all the experimental results must feature forward-path velocity commands corrupted by stationary noise. The results of this work provide behavioral support for the hypothesis that humans plan the motion components of their actions in terms of velocity.

## Introduction

Quite aside from its obvious scientific importance, a quantitative understanding of how neural activity in the central nervous system (CNS) manages the coordination of perception and action would have considerable practical value. Beyond the diagnosis and treatment of neurological disorders lies the promise of brain-machine interfaces—new ways to restore human capability after impairment or to augment it before injury. Already some remarkable advances have been reported (Carmena et al. 2003; Hatsopoulos and Donoghue 2009; Shoham et al. 2005). Notably, two successful demonstrations of controlling multi-joint robotic/prosthetic arms (with up to 22 degrees of freedom) interpreted the electrical signals recorded from the cerebral cortex as specifying velocity (Collinger et al. 2013; Hochberg et al. 2012). These studies reinforced compelling earlier demonstrations that complex drawing motions could be extracted from recordings of cortical neural activity that were interpreted as velocity commands in Schwartz et al. (Inoue et al. 2018; Moran and Schwartz 1999; Schwartz 1993, 1994).

Stochastic variation or ‘noise’ is ubiquitous in the neuromotor system (and in measurements of neural activity). The behavioral consequences of that noise depend sensitively on the neuromotor control architecture and how the noise is expressed in the neuro-mechanical system. The contribution of this paper is to report behavioral observations of the stochastic processes in human motor control and articulate model classes that can and cannot account for this behavior.

### Why command velocity?

What might be the advantages of velocity commands? Engineering may provide insight:

One example comes from amputation prosthetic applications. For practical reasons (especially their limited acceptable weight and volume) typical myoelectrically-controlled upper-limb prostheses are actuated by small electric motors with high-ratio speed reduction, which amplifies output torque in proportion, enabling lighter, more compact designs (Laffranchi et al. 2020; Psyonic 2023; Santina et al. 2018). The result is a non-back-drivable prosthesis quite different from natural limb behavior—external forces have minimal influence on prosthesis motion. By mapping myoelectric activity to (commanded) motor speed, maintaining a fixed posture (e.g. to carry a coat) is achieved by relaxing the controlling muscles. Fixed posture becomes a ‘default’ mode requiring no active intervention (and no unnecessary drain on batteries).

Another example is found in robot control. A typical task involves the motion of the robot end-effector (i.e. ‘hand’) but that motion is generated by actuators located at the robot’s degrees of freedom (i.e. ‘joints’). Joint motion uniquely defines task motion. However, execution requires mapping the planned task-space motion back to the corresponding joint-space motion. This requires inverting the kinematic relation between joint motion and task motion but that inversion often has no ‘closed-form’ representation. A practical alternative is to map task-space velocity to joint-space velocity (Whitney 1969). This linear algebra calculation uses the Jacobian matrix of the kinematic map. While the inverse Jacobian may not be unique, it is easy to add conditions that yield a well-defined solution (Murray et al. 1994).

Yet another example is found in aeronautic and aerospace applications. Flying requires control in six dimensions (three rotations and three translations). Among these dimensions, rotations provide a non-trivial challenge; a means to stabilize rotational motions was the essence of the Wright Brothers’ 1906 U.S. patent (Wright and Wright 1906). Finite angular rotations do not commute; the order in which they are performed affects a body’s resulting orientation. Unlike finite translations, they cannot simply be added as vectors. Remarkably, angular velocities are quite different: they commute (the order in which they are performed does not matter) and they may be added (linearly superimposed) as vectors. In practice, the most preferred representation of rotation for aircraft and spacecraft is via angular velocity relative to a body-fixed reference frame (roll, pitch, and yaw).

Of course, there is no guarantee that these engineering considerations apply to biology. Nevertheless, biology is subject to the same mechanical physics. If commanding velocity is a practical way to ‘work around’ the constraints of mechanical physics, we may expect related strategies in biology.

### Consequences of ubiquitous ‘noise’

Another important feature of the neuromotor system is the pervasive presence of stochasticity or ‘noise’ (Faisal et al. 2008; Harris and Wolpert 1998). Stochasticity is ubiquitous in both neurons and muscles.

Electroencephalographic (EEG) and electromyographic (EMG) data are notoriously noisy. While some of this noise is due to limitations of the measurement technology, the dominant component is due to intrinsically stochastic neuro-muscular behavior, phenomena ranging from the timing of action potentials to the dynamics of ion channels (Faisal et al. 2008). The variable that noise corrupts can have profound implications. In this work, we consider the behavioral consequences of velocity-level commands corrupted by stationary noise. If the central nervous system commands motion in terms of velocity, and those commands are affected by stationary noise, an immediate consequence is that noise on position would necessarily be non-stationary. For example, if the noise corrupting velocity commands is assumed to be white with constant strength (a common modeling assumption) the noise corrupting position would necessarily be Brownian (Wiener 1976), a ‘random walk’ phenomenon characterized by variance (second central moment) that grows linearly with time (Einstein 1905) and power spectral density declining in inverse proportion to frequency (-20 dB/decade on a log-log Bode magnitude plot).

To test this prediction would require experimental observation of unbounded variation; how might that be possible? Though it may seem counterintuitive, despite the manifestly bounded workspace of the human limbs, it is possible to experience unbounded behavior i.e., “the possibility of a finite and yet unbounded space” (Einstein 1916). Turning a crank is a simple example: a crank-and-pedal arrangement is used in a typical bicycle; hand cranks are common in manually-operated machinery. While the foot (respectively hand) is confined to a finitely bounded range of distances from a stationary pelvis (respectively thorax) the angular displacement of the bicycle crank (respectively hand crank) is unbounded; in principle, infinite angular displacement is possible. In the following we report observations of apparently unbounded variance in the human behavior of turning a simple hand crank.

Conversely, there are situations in which unbounded behavior cannot be observed experimentally. Upright posture provides an example; to maintain stable quiet standing, the center of pressure (CoP) of foot-floor interaction forces must remain within the base of support defined by the feet. However, that physical bound does not require the noise to be stationary; CoP variance may grow or decline with time—i.e. ‘drift’ as in a random walk—provided it remains within the base of support. The literature on human postural behavior confirms the presence of drift in quiet standing: the CoP of foot-floor interaction meanders within the foot-defined base of support in a manner resembling a random walk (Collins and de Luca 1993, 1995). Several descriptive models have been proposed to account for the ‘random walk’ nature of CoP variation (Collins and de Luca 1993, 1995; Delignières et al. 2011; Kuznetsov et al. 2013; Peterka 2000). Here we show that this non-stationary behavior can emerge in three distinct ways. We present new experimental evidence and show that only one of these three models is plausibly competent to account for all of the experimental evidence.

## Results

### Crank Turning

A crank-turning experiment was conducted with 10 subjects (Fig 1.a). Crank turning was selected since it is particularly common in human activities of daily living (Petrich et al. 2022) but has the remarkable property of allowing unbounded motion despite the finite workspace of the human arm. Subjects were instructed to turn a crank at a constant speed in the clock-wise direction. Visual feedback of the rotational speed was provided. No explicit display of the angular position of the crank was provided, though normal proprioceptive feedback was available.

**Fig 1.**
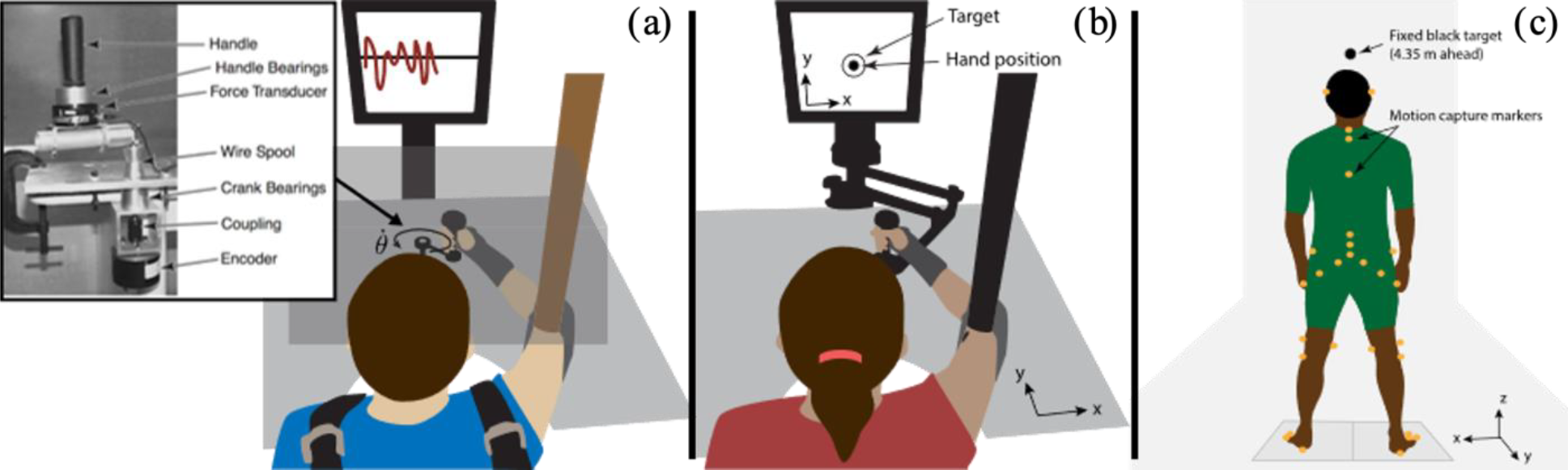
Experimental setups. (a) Crank turning: the crank displayed in the inset was used to provide a circular constraint. Vision of the arm and crank was occluded but the subject was provided with visual speed feedback. The wrist was braced, the upper arm was supported by a sling, and the shoulders were strapped to a chair. (b)Hand posture: the InMotion2 robot was used to measure hand position. Visual feedback of the hand position was provided on a display. Vision of the arm and the robot was not occluded. The arm was supported by a sling. (c)Quiet standing (dos Santos et al. 2017): subjects stood upright with their arms by their side, with their gaze fixed on a black target on a wall 4.35 m ahead. Motion capture markers were placed on their body. Further details of the three experiments are provided in the Material and Methods section.

The experimental results for a single subject performing the crank-turning task are presented in Fig 2. The three panels show: (a) the time trajectories of the crank angle over multiple crank rotations, (b) the crank-angle variance, and (c) the Bode magnitude plot of the related PSD. The variance was computed across the ensemble (21 trials), while the PSD was computed for each trial and then averaged across trials.

**Fig 2.**
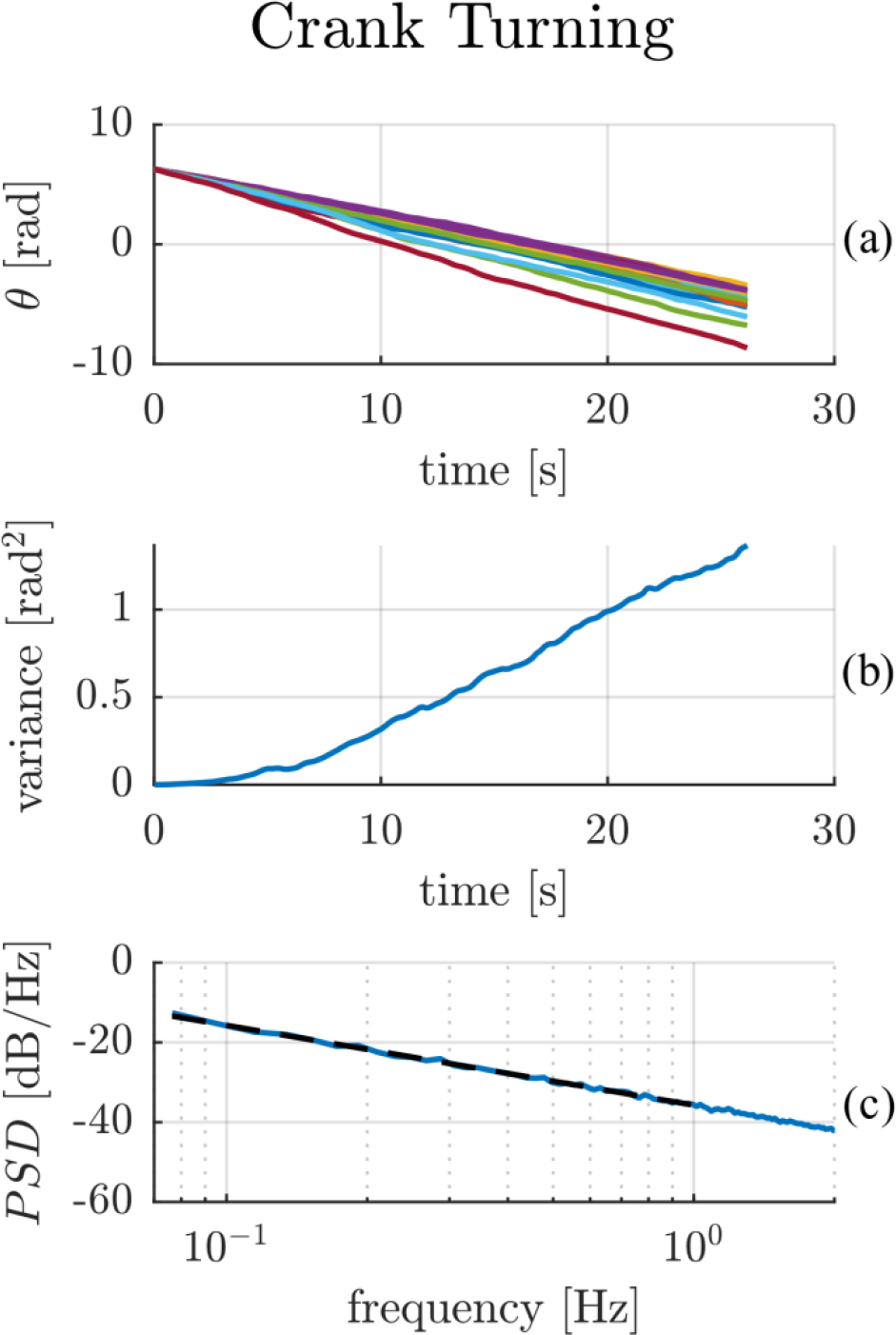
Crank turning experimental results for a single subject. The top plot (2.a) shows the crank angular position trajectory over time for multiple trials. The central plot (2.b) presents the position variance computed across trials. The bottom plot (2.c) shows a Bode magnitude plot of the average power spectral density of the angular position (solid blue line) compared to the best-fit line with a slope of -20.01 dB/dec (black dashed line).

The single-subject data in Fig 2 exhibit Brownian behavior – an unbounded, linearly-growing variance of crank angle with time 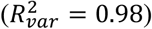. In the frequency domain, we expected a −20 dB/dec slope in the Bode magnitude plot of the crank angle, which we observed (−20.01 *dB/dec*,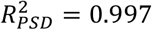), with 95% CI [−20.35 − 19.67] dB/dec at the lowest frequencies (0.07-1 Hz).

The same behavior was observed across all subjects, with an average 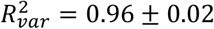 for the variance slope, and an average Bode magnitude slope of −20.89 dB/dec 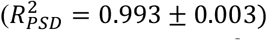 with 95% CI of [−21.40 − 20.36] dB/dec at low frequencies. The Supplementary Information contains 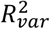and 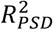for the variance and PSD models and the 95% confidence intervals of the Bode magnitude slopes for all subjects.

### Hand Posture

A postural task for the upper limb was devised (Fig 1.b) to test the presence of bounded Brownian-like behavior in a static positioning task. 10 subjects were asked to hold their hand at a fixed position for 4 minutes while visual feedback of hand position was provided. A sling supported the arm to minimize the gravitational load, while a robotic manipulandum with negligible friction was used to measure the planar (x,y) trajectory of the hand. 10 trials were performed.

The experimental results for a single subject performing the hand-posture task are presented in Fig 3. The three panels show: (a) the time trajectories of hand position over the 10 trials, (b) the hand-position variance, and (c) the Bode magnitude plot of the PSD of hand position. The x-coordinate of the hand position is presented; the y-coordinate (omitted for clarity) exhibited similar behavior.

**Fig 3.**
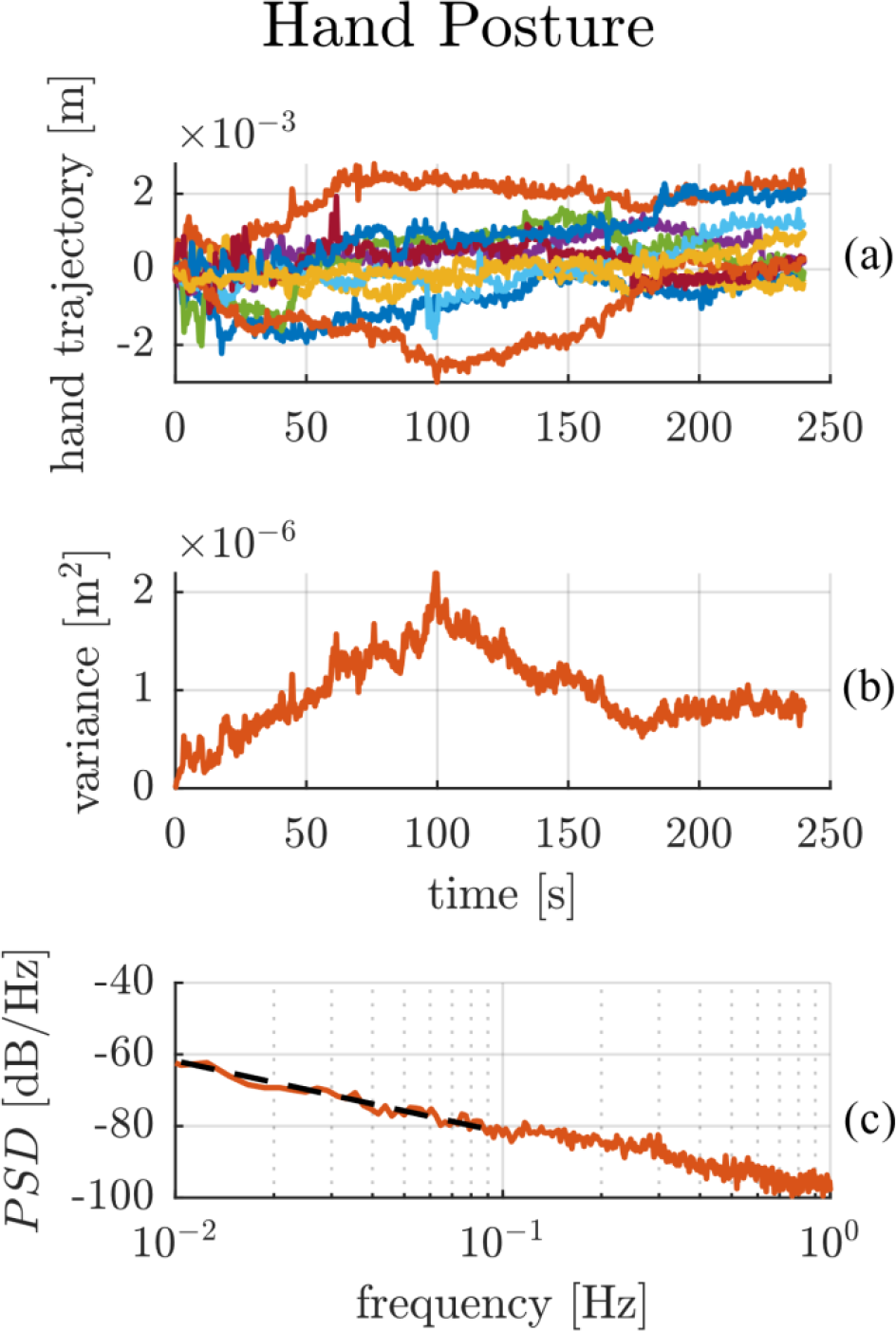
Hand posture experimental results of a single subject. The top plot (3.a) shows the trajectories of the ‘x’ Cartesian coordinate of the hand over time for multiple trials. The central plot (3.b) presents the position variance computed across trials for the ‘x’ direction. Note the obviously non-stationary character of the noise process. The bottom plot (3.c) shows a Bode magnitude plot of the average power spectral density of hand position in the ‘x’ direction (solid red line) compared to the best-fit line with a slope of -18.46 dB/dec (black dashed line).

Once again, the single-subject data show that hand-position variance grew approximately linearly with time over the first minute in both the ‘x’ and ‘y’ directions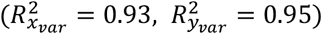, suggesting that the presence of a Brownian process is an intrinsic feature of human neuromotor behavior. However, unlike in the crank-turning task, the hand-position variance reached a ‘break point’ 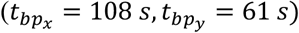 after which the variance stopped its linear increase. In fact, linear regression of the variance computed over the whole time period showed a poor fit with 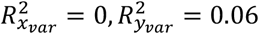. Over the low-frequency range (0.01-0.1 Hz) the Bode magnitude plot of the PSD displayed a −18.46 dB/dec slope 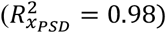, with 95% CI [−21.08 − 16.84] dB/dec in ‘x’ and a −21.39 dB/dec slope 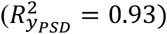 with 95% CI [−23.16 − 19.61] dB/dec in ‘y’.

In all 10 subjects, a consistent linearly-growing variance up to a break point was observed, with an average 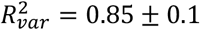. The break point between linear growth and bounded variance varied across subjects, ranging from a few seconds (1-5 s) to minutes (100-135 s). Subjects’ low-frequency PSD Bode magnitude plots presented good linear fits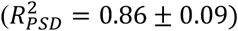, with an average best-fit slope of −16.69 dB/dec and an average 95% CI of [−18.48 − 14.90] dB/dec in both directions. The minimum frequency for the linear fit was 0.01 Hz for all subjects except subject no. 5 who had a minimum frequency for the linear fit of 0.04 Hz. The flattening of the low-frequency Bode magnitude slope may be attributed to the boundedness of the process, and was reproduced in simulation (see the Descriptive Model below). The Supplementary Information reports the variance and coefficients of determination 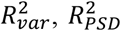and breakpoints, as well as the average PSD Bode magnitude slope and related 95% confidence intervals for all 10 subjects.

### Quiet Standing

Upright standing posture has been a prevalent behavioral paradigm in the study of the stochastic processes underlying human motor control. To enable a comparison between the upper limb experiment presented above with behavior during natural upright posture, the same analysis was run on the experimental data collected by dos Santos et al. (dos Santos et al. 2017). This database contains observations of 49 subjects standing for 60 seconds at a time (Fig 1 right panel). Motion-capture markers were placed on the body to determine the center of mass (CoM) trajectory and joint angles over time.

In our study, the CoM trajectory in Cartesian coordinates during the normal quiet-standing trials (eyes open, standing on a flat rigid surface) for 26 young unimpaired subjects was considered. The CoM was chosen as a macroscopic representation of overall body behavior; it may be considered equivalent to the end-effector position of an open-kinematic-chain model of upright human posture.

The experimental results for a single subject are presented in Fig 4. The two panels show: (a) the time trajectories of the CoM position over the 3 trials, and (b) the related Bode magnitude plot of the PSD. The x-coordinate of the CoM is presented; the y-coordinate (omitted for clarity) exhibited similar behavior.

**Fig 4.**
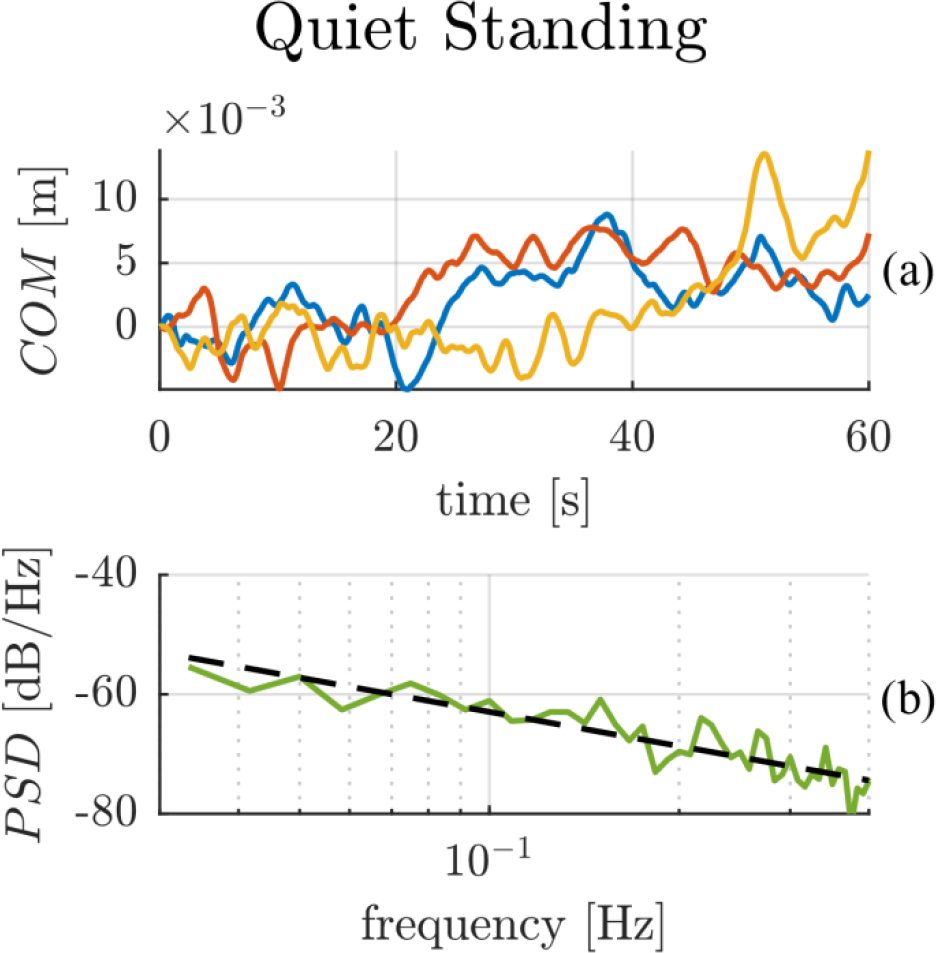
Quiet standing experimental results for a single subject. The top plot (4.a) shows the ‘x’ component of the center of mass trajectories over time for multiple trials. The bottom plot (4.b) presents the average Bode magnitude plot of the power spectral density of the ‘x’ component of the center of mass trajectories (solid green line), while the black dashed line presents the best-fit line with slope -19.05 dB/dec.

The CoM trajectory of the subject (Fig 4.a) shows evidence of random fluctuations. The Bode magnitude plot of the PSD of these fluctuations follows a linear 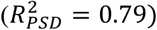 slope of −19.05 dB/dec in ‘x’, with 95% CI [−22.09 − 16.00] dB/dec, and a slope of −18.27 dB/dec 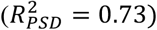 in ‘y’ with 95% CI [−21.69 − 14.86] dB/dec, over the low-frequency range (0.03-0.4 Hz) as shown in Fig 4.b.

A similar behavior was observed across all subjects, with an average Bode magnitude slope of −16.06 dB/dec 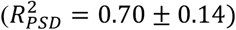 with 95% CI on both directions of [−19.12 − 13.00] dB/dec, displaying once again a flatter-than-Brownian slope. The Supplementary Information contains the best-fit PSD slopes, 95% confidence intervals, and coefficients of determination *R*^2^ for all subjects.

### Descriptive Model

The crank-turning experiments displayed the signature of a purely Brownian process, while the hand-posture experiments showed signs of variance growing linearly with time only up to a certain time point, after which the variance stopped increasing. These observations are consistent with our expectations: position variance must be bounded for postural tasks. However, it is possible that all three tasks share a common control architecture that produces a Brownian process, with an additional layer of control responsible for bounding the variance in postural tasks. Here, we develop a descriptive model assuming that a similar control structure underlies all three tasks.

A general description of all three tasks requires a model with closed-loop negative feedback. In this case, Brownian behavior, in position, can emerge from only three sources: (i) an additive external disturbance as originally proposed by Peterka (Peterka 2000); (ii) a free integrator in the closed-loop dynamics; and (iii) a forward-path (reference) velocity command, corrupted by stationary noise and integrated to produce a position reference command. Figure 5 provides a schematic representation of these three possibilities. A detailed description of all three models is presented in the Supplementary Information. In the following we assess their biological plausibility.

**Fig 5.**
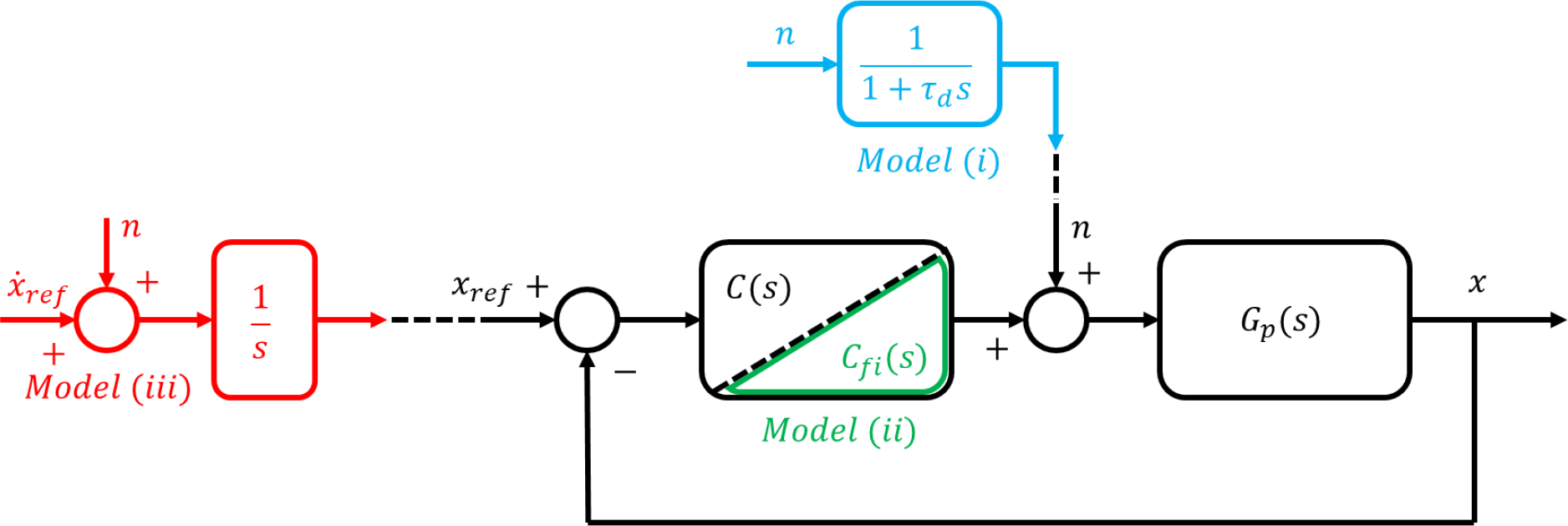
Block diagram of a closed-loop negative feedback control system with the three different ways to generate Brownian-like behavior in position: model ‘i’ (light blue) consists of an additive external stationary noise ‘*n*’ processed through a first-order low-pass filter (a ‘leaky integrator’); model ‘ii’ (green) requires a constrained design of the controller ‘*C*_*fi*_ (*s*)’ such that the closed-loop dynamics exhibits a free integrator; model ‘iii’ (red) introduces a velocity level command 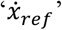 affected by stationary noise ‘*n*’ and integrates it to obtain the reference position command ‘*x*_*ref*_’. ‘*G*_*p*_(*s*)’ represents the open-loop dynamics of the physical system being controlled, whether it is the arm for the crank-turning and hand-posture tasks, or the full body in upright stance for the quiet standing task. ‘*G*_*p*_(*s*)’ receives joint torque commands as input, and outputs the position measurement of interest for the given task.

### Additive external disturbance – Model ‘i’

To account for the boundedness of postural tasks (hand posture and quiet standing) the first model (Peterka 2000) assumes stationary noise modified by a first-order low-pass filter (a ‘leaky integrator’) added to the input to the physical system (e.g. muscle forces applied to the skeleton). To reproduce the observed bounded postural behavior, this model requires the process time-constant (τ_*d*_ in Fig 5) to be fast enough that the bound is reached in a finite time window as in typical observations (see experimental behavior of Fig 3.a). At the same time, the process time-constant τ_*d*_ must be slow enough to reproduce the apparently unbounded variance of the crank-turning data in Fig 2. Strictly Brownian behavior over the time window of our crank-turning observations (26 s) would require a time-constant of about 75 s, which seems implausibly long. Based on the data we observed, these two requirements appear to be incompatible.

### Free integrator in the closed-loop dynamics – Model ‘ii’

The second model requires the implementation of a controller capable of producing free-integrator behavior in the closed-loop dynamics. From a classical control perspective, this means placing a pole at the origin of the closed-loop transfer function and that requires complete knowledge of the system to be controlled (‘*G*_*p*_(*s*)’ in Fig 5). In the context of standing balance, the extrinsic de-stabilizing gravitational load must be exactly balanced by the intrinsic neuro-muscular stiffness, a topic debated in the standing-balance literature, which makes this model less plausible.

A free integrator would guarantee a strictly Brownian behavior in position, thus reproducing the crank turning data of (Fig 2). To account for the postural data (Figs. 3 and 4), some process to limit position variance is required in addition.

### Forward-path (reference) velocity command, corrupted by stationary noise – Model ‘iii’

The third model guarantees the emergence of Brownian postural behavior by means of a forward-path velocity command 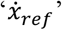 corrupted by stationary noise ‘*n*’ (see Fig 5). In this case a Brownian process is already embedded in the command signal ‘*x*_*ref*_’ and no particular knowledge of the control system (*C*(*s*) or *G*_*p*_(*s*)) is required. Similar to model ‘ii’ (the free integrator), some form of correcting action is required to prevent position variance from growing without bound during postural tasks.

### Intermittent control

One biologically-plausible means of bounding variance is through intermittent control actions triggered by crossing a threshold. Intermittent control has been proposed frequently in the motor neuroscience literature (Bottaro et al. 2005, 2008; Gawthrop et al. 2011; Loram et al. 2022, 2011, 2012; Markkula et al. 2018; Milton 2013; Morasso et al. 2020).

Of the three proposed models, we report here the simulation results of model ‘iii’ with intermittent control, which was able to account for all the experimental evidence. A total of 200 numerical simulations per condition were performed to assess the competence of the proposed model to reproduce the experimental observations. Fig 6 shows the model performance in the three different tasks: crank turning, hand posture, and quiet standing.

**Fig 6.**
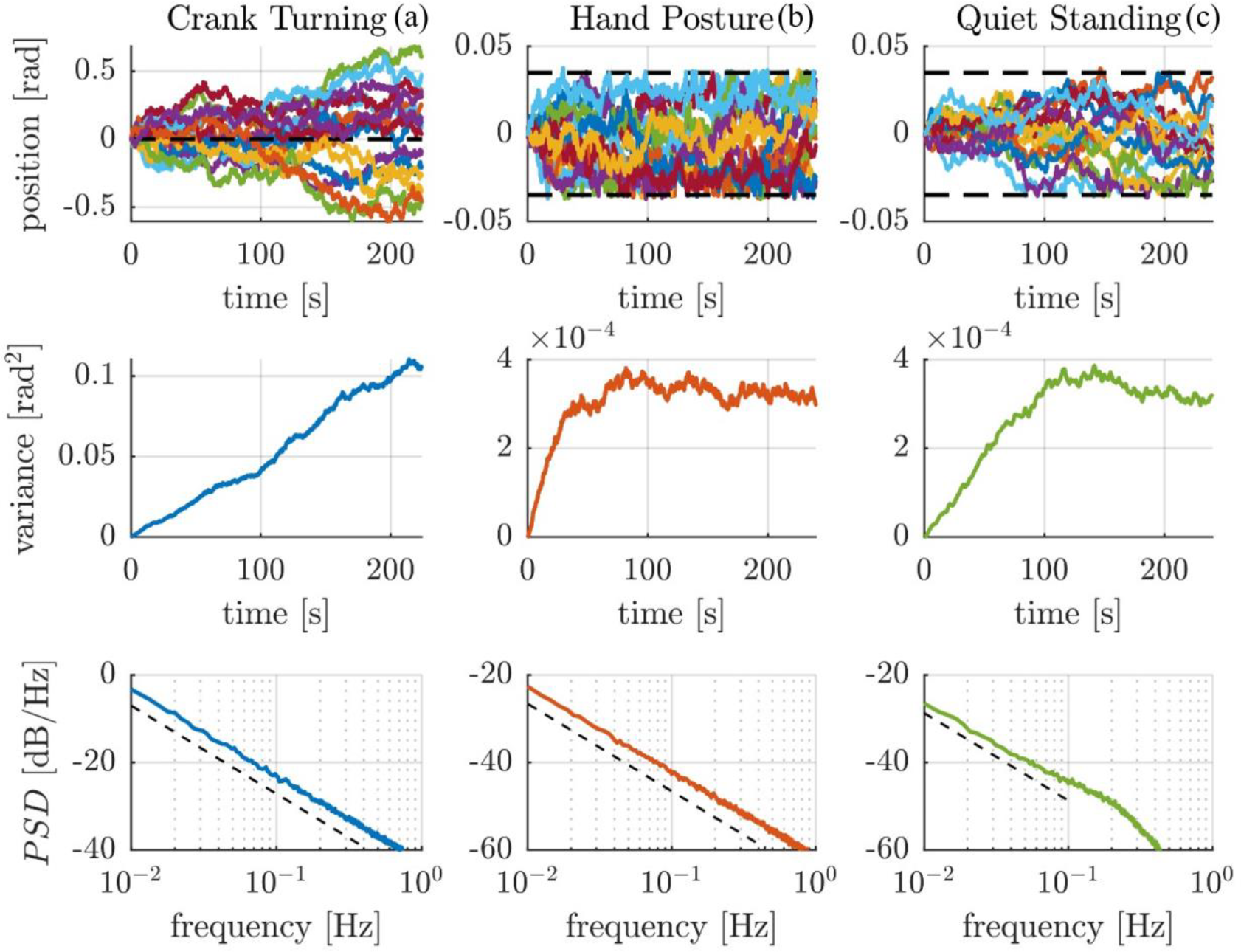
Simulation results of the proposed control model across three tasks: (a) crank turning, (b) hand posture, and (c) quiet standing. The top row of plots shows the time evolution (colored lines) of position relative to the expected nominal trajectory in the three cases; the black dashed line in the crank-turning trajectory plot represents the expected nominal trajectory for the target crank-turning speed, while the black dashed lines in the right two trajectory plots present the thresholds to trigger intermittent control action in the posture control tasks. The middle row of plots shows the variance over time across the ensemble. The bottom row presents the Bode magnitude plots of the average power spectral densities in the three different tasks. An additional -20 dB/dec dashed-black reference line is added to facilitate interpretation of the results.

The model reproduced both the unbounded Brownian behavior during crank turning (Fig 6.a) as well as the bounded behavior during hand posture and quiet standing (Figs. 6.a and 6.b). The PSDs (Fig 6) present a low-frequency (0.01 Hz – 1 Hz for crank turning and hand posture, 0.01 Hz – 0.1 Hz for quiet standing) Bode magnitude slope consistent with the experimental evidence: crank-turning: −19.70 dB/dec with 95% CI [−19.78 − 19.61] dB/dec; hand posture: −19.42 dB/dec with 95% CI [−19.50 − 19.34] dB/dec; and quiet-standing: −17.94 dB/dec with 95% CI [−18.35 − 17.53] dB/dec.

## Discussion

A Brownian process – also known as a Wiener process or random walk – is a non-stationary process characterized by variance that grows linearly with time (Einstein 1905). First discovered by botanist Robert Brown in 1828, it found a more formal definition in one of Einstein’s *annus mirabilis* papers in 1905 (Einstein 1905), and was rigorously described mathematically as a stochastic process by Norbert Wiener many years later (Wiener 1976). Evidence of Brownian processes has been found in many fields: the behavior of particles in a stationary liquid, in chemistry (Kramers 1940), electromagnetism (Kurşunoğlu 1962), fluid-dynamics (Hauge and Martin-Löf 1973), and the trends of stock prices (Osborne 1959). Here we provide evidence that it is a ubiquitous (and perhaps essential) feature of human motor behavior.

### Experiments

The experimental PSDs in the crank-turning case show a slope remarkably close to −20 dB/dec (Fig 2.c). This behavior indicates a strictly Brownian process at low frequencies. In the hand-posture and quiet-standing cases, we observed a flattening of the PSD slope at low frequencies (Figs 3.c and 4.b). At higher frequencies, the slope varied.

Reaching with the upper limb has been studied widely in motor neuroscience (Abend et al. 1982; Burdet et al. 2001; Flash and Hogan 1985; Gulletta et al. 2020; Won and Hogan 1995). A kinematic constraint provides an intermediate stage between unconstrained (free) motion and interaction with complex dynamics. Circularly constrained motion, such as crank-turning, has been less widely studied (Hermus et al. 2023; Ohta et al. 2004; Russell and Hogan 1989), despite being ubiquitous in everyday manipulation (e.g., turning a steering wheel or opening a door). In fact, opening a door was reported to be the most common activity of daily living (Petrich et al. 2022). The crank-turning task occupies a finite region of the arm’s workspace; as a result, the displacement of the hand relative to the thorax is bounded. However, the angular position of the crank is unbounded. Thus, the variance of angular position may grow without bound, and in fact it grew linearly (Fig 2.2), indicating a Brownian process. It is worth emphasizing that this particular task allowed an unbounded task-space motion to be executed by motor actions in a finite joint space.

In the postural tasks, such as maintaining a static hand posture or quietly standing, the results also matched the predictions. In these cases, a Brownian-like behavior was still observed as a growing position variance until it reached a limit (Fig 3.b). This was confirmed by the position trajectories (Fig 3.a) where variance initially grew but was ultimately bounded. All subjects exhibited a transition between approximately linear growth and boundedness, but its onset varied among subjects, ranging from a few seconds to hundreds. Interestingly, a later onset of the bounded behavior corresponded to PSD slopes closer to an unbounded Brownian process (−20 dB/dec), while an earlier onset correlated with flatter PSD slopes.

The existing literature has reported phenomena similar to Brownian motion, often termed “drift” or “sway”, in the CoP trajectory during quiet standing. The origin of this “drift” and the underlying human motor control mechanisms have been discussed at length. Collins and De Luca were the first to suggest the presence of “fractional Brownian” behavior in CoP postural sway, proposing an alternation between “do-nothing” (open-loop) and “do-something” (closed-loop) control actions to explain the observations (Collins and de Luca 1993). Thereafter, several studies considered many aspects of this postural “sway”: its dependency on sensory information (Collins and de Luca 1995), its possible decomposition into different motor control mechanisms (Zatsiorsky and Duarte 1999), the possibility of describing it with linear models (Peterka 2000) or intermittent control (Loram et al. 2022; Morasso et al. 2020), the kind of stability necessary to explain it (Bottaro et al. 2005, 2008), or the use of different mathematical methods to characterize it (Delignières et al. 2011; Hernandez et al. 2016; Kuznetsov et al. 2013). Similar studies were extended to other tasks involving balancing or pointing objects with the upper limbs (Cabrera and Milton 2002; Goh et al. 2015). In these cases, too, related work studied, for example, the influence of sensory information (Brown et al. 2003) or the underlying control mechanisms (Cabrera and Milton 2002).

A fractional Brownian process is associated with chaotic dynamics and implies a nonlinear growth of variance with time, unlike the linear growth of variance that characterizes a Brownian process. Our results, however, indicate no need to postulate fractional behavior. In fact, we believe that the apparent fractional behavior reported in previous work (Collins and de Luca 1995; Delignières et al. 2011; Kuznetsov et al. 2013) may be a consequence of the inertial properties of the system and the bounded behavior during the postural tasks. A detailed example is presented in the Supplementary Information.

The results reported here establish that a Brownian process is a good description of intrinsic position variability. However, some additional process must act to limit position variance in postural tasks. One possible explanation can be found in the theory of intermittent feedback control. Intermittency in human motor actions has been reported since Woodworth in 1899 (Woodworth 1899), and fundamentally arises – in its simplest form – by alternating a “do-nothing” region for small feedback errors within which there is no control action, with “do-something” regions to correct for larger feedback errors. Several studies have supported intermittent control in human motor behavior (Bottaro et al. 2005, 2008; Cabrera and Milton 2002; Collins and de Luca 1993; Craik 1947; Gawthrop et al. 2011; Markkula et al. 2018; Morasso et al. 2020).

## Models

The proposed descriptive model (model ‘iii’ of Fig 5) exhibits Brownian behavior by means of stationary noise added to a reference (forward-path) velocity command. This is equivalent to including a “free integrator” upstream of the interactive dynamics in a Norton equivalent network (Hogan 2017). To bound the variance in postural tasks, an intermittent controller was included in the forward path where the “do-something” region outside the feedback-error thresholds generated a corrective action as a velocity command. For simplicity, linear representations of the open-loop system dynamics *G*_*p*_(*s*) and compensator *C*(*s*) were assumed but that is not essential. We emphasize that this work identified a feature in one of the lowest levels of a complex control system. It is silent about the form of the higher-level controller as long as it presents intermittent feedforward action as velocity commands. Interestingly, this is consistent with recent robotic implementations (Hwangbo et al. 2019). Fig 6 shows that this model can reproduce all of our experimental observations: crank turning, hand posture, and quiet standing.

Model ‘iii’ is not the only possible way to generate a Brownian process in position: there are two alternatives. The first alternative (model ‘i’) was inspired by Peterka (Peterka 2000) who showed that the phenomena reported by Collins and De Luca (Collins and de Luca 1993) could be reproduced by a continuous linear time-invariant controller with a low-pass-filtered stationary noise disturbance, with no need for intermittent ‘open-loop’ and ‘closed-loop’ action. Peterka’s model requires an appropriate tuning of the time constant (‘τ_*d*_’ of Figure 5) to produce the desired bounded behavior in postural tasks. This time constant ranged between 10 s to 120 s in Peterka’s work, which appears difficult to justify for human motor control. In our simulations, we could reproduce the bounded behavior of hand posture and quiet standing with the time constant ranging from 20 s to 160 s. However, in order to reproduce the crank turning behavior we had to use a time constant of at least 78 s and as much as 240 s. As this model requires such a wide range of time constants to account for all of the experimental evidence, we believe that it is not the most suitable to describe all three tasks investigated in this work.

The second alternative model (‘ii’) requires the system’s closed-loop transfer function to exhibit exactly one pole at zero to produce a Brownian process in position. In principle, this could be achieved by a controller that compensates exactly for gravitational destabilization during upright posture or by nullifying arm stiffness during gravity-neutral hand posture. Whether intrinsic joint stiffness is sufficient to stabilize standing posture has been debated in the balance literature (Loram and Lakie 2002; Winter et al. 2001), and exactly balancing extrinsic and intrinsic loads seems challenging, to say the least. However, to cancel gravitational stiffness requires less joint stiffness than to stabilize posture and it might be interesting to re-consider this matter in light of this reduced requirement. To maintain a gravity-neutral posture of the hand would require imposing a negative stiffness to compensate for intrinsic muscle impedance, which is positive (Bennett et al. 1992; Mussa-Ivaldi et al. 1985). Feedback processes (e.g. via spinal reflexes) could generate ‘negative stiffness’, and apparently do so in some circumstances (Gillard et al. 1999, 2000). In this second alternative model, an intermittent control action would suffice to bound the variance in postural tasks. Nonetheless, this model fails to reproduce the unbounded task i.e., crank-turning. Having a net stiffness equivalent to zero is equivalent of having a velocity tracking controller with no integral action (stiffness) which will cause a steady-state error in the velocity tracking performance, and thus position – integral of velocity – will grow with a different slope (rate) compared to the required reference. This is not what we observed experimentally i.e., the measured position grew with the same slope (rate) of the reference target velocity.

Of course, it is possible that different control architectures underlie different motor tasks, and one may argue against constraining ourselves to a model that can describe all of the experimental observations in the three different tasks. However, given the pervasiveness of drift, Brownian behavior appears to be a fundamental feature of neuromotor control, and it is plausible that this Brownian process is a manifestation of some common control architecture shared across the various motor tasks, i.e. velocity-level planning.

Which mechanical variables best correlate with neural activity is controversial, and irrelevant in some schools of thought. This is probably because many factors are important to successful motor behaviors. Limb velocity, displacement, force, and force rate initially presented plausible correlation values (Humphrey et al. 1970). Knowledge of neural encoding is still evolving. More recently, another perspective has emerged, suggesting that muscle activation may result from the collective activation of many neurons which generate rotational dynamics to produce a learned feedforward action that plays out over a finite duration (Churchland et al. 2006; Shenoy et al. 2013). However, our discussion of velocity commands does not preclude its relevance to such a representation.

Apart from successful implementation in brain-machine interfaces, the importance of velocity commands in motor tasks has been highlighted in numerous other studies. Evidence of velocity-level planning is found in the work of Atkeson and Hollerbach (Atkeson and Hollerbach 1985) on spontaneous reaching tasks, where they observed highly-stereotyped speed profiles despite substantial variation of hand paths. Krebs et al. reported evidence of stereotyped upper-limb speed profiles in the earliest spontaneous movements of recovering stroke-survivors (Krebs et al. 1999). In a study of re-targeting reaching movements, Flash and Henis found results consistent with stereotyped hand speed profiles (Flash and Henis 1991). Evidence of velocity commands has also been found in other neural circuits: Robinson, studying saccadic eye movements, proposed that premotor neurons encoded movement velocity and duration (Robinson 1970, 1973); Rodman and Albright reported velocity coding in the middle temporal visual area cells of macaque (Rodman and Albright 1987); Keshavarzi et al. recently reported that retrosplenial cortical neurons reliably track direction and speed of head turning (Keshavarzi et al. 2022). All these studies point to velocity as a significant variable in neuromotor control, perhaps for the same reasons that velocity commands are useful in engineering applications.

The observations of Brownian processes in the human motor tasks presented in this work further support the idea of velocity-level planning for movement control: a model with forward-path velocity commands corrupted by stationary noise, and intermittent control action during postural tasks, is capable of reproducing all of our experimental observations. If the brain interprets descending (forward-path) motor commands in terms of velocity (Moran and Schwartz 1999; Schwartz 1993, 1994), then because of noise, the resulting position must express a Brownian behavior, at least before intermittent correction—just as we observed. To the best of our knowledge, this model, together with the experimental evidence presented, provides behavioral support for the hypothesis that some aspects of neural activity can be interpreted as velocity commands.

## Material and Methods

### Subjects

Two different human subject experiments were conducted at the Massachusetts Institute of Technology: crank turning (10 right-handed college-aged male subjects), and static hand posture control (4 males and 6 females, ages 23-36). All participants gave informed, written consent before the experiment. The informed consent and experimental protocols were reviewed and approved by the Institutional Review Board for the Massachusetts Institute of Technology. The quiet standing data came from a public data set of experiments conducted by Santos et al. (2017) and consisted of data from 49 subjects (dos Santos et al. 2017); the data for the 26 young unimpaired subjects in the study (15 males and 11 females, ages 18-40) were used in this paper.

### Crank Turning Experiments

Subjects turned a crank mounted on a high precision incremental optical encoder/interpolator set (Gurley Precision Instruments encoder #8335-11250-CBQA, interpolator #HR2-80QA-BRD) with a resolution of 0.0004 degrees per count.

During the experiment, the subject’s arm was occluded from view by a wooden structure, which did not limit the range of motion. The upper arm was suspended by a canvas sling connected to the ceiling using a steel cable; the upper arm and lower arm were in the plane of the crank. The subject sat in a chair with a rigid back, while the shoulder was restrained by a harness attached to the back of the chair. The subject was positioned such that the crank, with radius 10.29 cm, was well within the workspace of the arm. Fig 1 shows a graphical representation of the experimental setup.

The encoder, sampling at 200 Hz, was connected to a set of counters and to the computer via digital I/O. The visual display, also generated by the computer, was on a 17-inch monitor (311 x 238 mm, resolution 1280 x 1024, 76 Hz) which was mounted approximately 75 cm from the subject’s eyes. The experiment was divided into two unequal sections: 2 blocks of trials at the subject’s preferred or ‘comfortable’ speed and 6 blocks of trials at a visually-instructed speed.

The data were reported by Hermus et al. (Hermus et al. 2020). The results presented herein only include analysis of the slow, clockwise-turning condition. However, details of the entire experimental procedure are included for completeness.

At the start of the experiment, subjects performed 20 trials at their preferred speed, 10 trials in the clockwise direction (CW) and 10 in the counterclockwise direction (CCW); both conditions were blocked, in random sequence for each subject; each trial lasted 8 seconds. Subjects were not provided with any visual feedback during these trials. Thereafter, subjects performed 6 blocks of 30 trials, each with visual specification of 1 of 3 target speeds, in either CW or CCW directions. The order of the speed and direction blocks was pseudo-randomized across subjects. The three speeds were selected to cover a significant range: 0.075 rev/s was extremely slow (required over 13 s per revolution), 0.5 rev/s was close to subjects’ preferred speed, and 2.0 rev/s was close to the fastest speed that subjects could turn the crank. Visual feedback on the monitor displayed the target speed, as well as the subject’s real-time hand speed; the horizontal axis was time, and the vertical axis was speed. Target speed was displayed as a continuous horizontal line in the middle of the screen. The width of the screen corresponded to the time of the trial. 7 catch trials without visual feedback were included in each block, leaving 23 trials per block with visual feedback.

In the slow-speed conditions, each trial lasted 45 s. This yielded about 3.4 turns of the crank for the slow condition. The duration of the slow-speed trials was chosen as a compromise between acquiring adequate data and avoiding subject fatigue. The first and last cycle were removed to eliminate transients. This yielded a 26-second trial duration for analysis. The first 2 trials of each block were considered to be familiarization trials and were thus removed together with the 7 catch trials without visual feedback, leaving 21 trials per subject for analysis.

### Hand Posture Experiments

In the hand posture experiment, subjects were asked to hold the handle of an InMotion2 robot (Interactive Motion Technologies Inc.) and maintain a static hand posture at the center of a circle of radius 2.5 cm. The InMotion2 is a highly back-drivable two-link manipulandum. In the experiments, the robot served only as a position measurement device as the actuators were open-circuited. InMotion joint positions were measured by a 16-bit/rev encoder at ∼350 Hz using a Compact RIO real-time processor. Data were interpolated to a sampling rate of 200 Hz.

The experimental setup was the same as the previous crank experiment except that the crank was replaced with the InMotion2 robot, vision of the hand was not occluded, and the visual feedback was changed to show the fixed target position (given by a circle of radius 2.5 cm) and the subject’s actual hand position (given by a dot of radius 1 cm). Fig 1 shows the experimental configuration. Each trial lasted 4 minutes, and was repeated 10 times for each subject, with breaks between trials.

### Quiet Standing Experiments

The quiet standing experiments were conducted by Santos et al. (2017) in the Laboratory of Biomechanics and Motor Control at the Federal University of ABC, Brazil (dos Santos et al. 2017). Subjects were asked to stand barefoot, as still as possible, with their arms by their sides, under four conditions: on a rigid surface with eyes open, on a rigid surface with eyes closed, on a compliant surface (foam blocks) with eyes open, and on a compliant surface with eyes closed. Trials lasted 60 s and were performed 3 times per condition in random order. 3D kinematics were measured using a motion capture system, with data sampled at 100 Hz; post-processing was performed to estimate the center of mass (CoM) trajectory in Cartesian coordinates, as well as joint angles. Fig 1 shows the experimental configuration. Further experimental details can be found in the article by Santos et al. (2017) (dos Santos et al. 2017).

The results presented herein include data for the first condition (eyes open, rigid surface), and for the 26 young unimpaired subjects. The estimated CoM trajectories included in the public data set were used in the data analysis. Note that the chosen (x,y,z) coordinate directions (see Fig 1) are different from those used in the public data set by Santos et al. This was done to have the (x,y) coordinates in the horizontal plane for both quiet standing and hand posture.

### Data Analysis

The data - crank angular position, hand position (x,y), and CoM position (x,y) - were processed using MATLAB (version 2022a). Three different quantities were analyzed: (i) the time trajectory of the position data over each trial, (ii) their variance-over-time computed across the ensemble, and (iii) their average power spectral density.

The time trajectory was directly extracted from each trial of the collected data. For all research studies (crank turning, hand posture, and quiet standing), data from all trials were aligned such that the initial data point was taken to be at the origin, namely 0 *rad* for the crank turning angle, and (0,0) *m* for the hand and CoM Cartesian coordinates.

The variance across the ensemble was computed at each time step (*t*_*s*_ = 0.005 *s*) using the time trajectories (aligned at the initial data point) with the following equation:

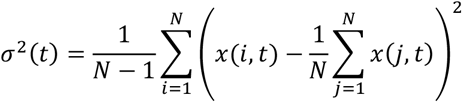

where *i* and *j* represent the trial indices ranging from 1 to N (number of trials), and *x*(*i, t*) represents the value of the considered variable at time *t* during the *i*-th trial.

The power spectral densities (PSDs) were computed for each trial with Welch’s method (Solomon, Jr, O M 1991) using each entire trial as one window of data (in order to obtain spectral estimates down to the lowest possible frequency) and averaging across trials. The data were pre-processed by removing the average crank angle trajectory from the crank angle data, and removing the mean position over each trial for the hand position and CoM position data. The PSDs were post-processed to remove the lowest-three frequency points, as these depended sensitively on the data processing technique used (see the Supplementary Information for details of the pre- and post-processing decisions).

### Modeling

The descriptive mathematical model, represented in the block diagram of Fig 7, was implemented in Simulink/MATLAB (v 2022a) to simulate the behavior of the three different motor tasks: crank turning, hand posture, and quiet standing. For each of these cases, a single-degree-of-freedom model was considered for simplicity.

**Fig 7.**
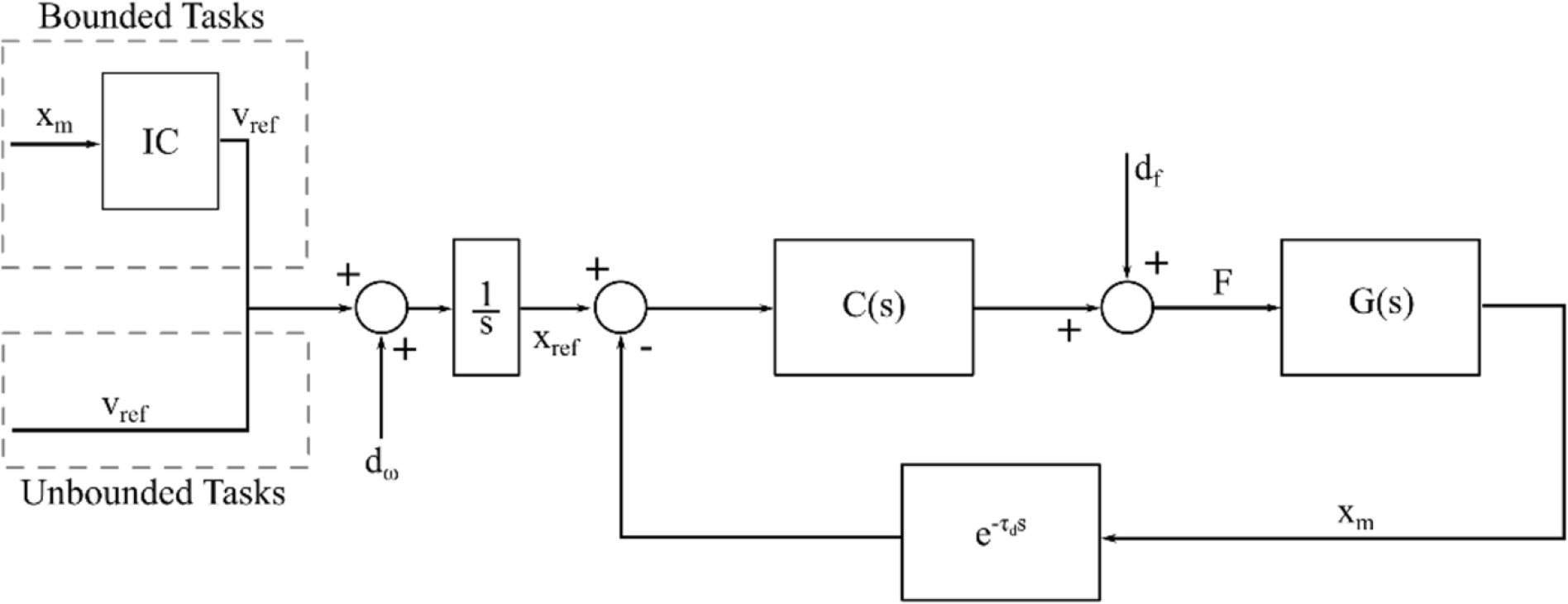
Block diagram of the modelled control scheme. G(s) represents the open-loop system dynamics, where s is the Laplace variable; C(s) is the closed-loop control action; IC represents the intermittent controller necessary to bound position variance; *d*_*f*_ and *d*_ω_ represent additive noise on the control action (force/torque) and on the reference velocity command, respectively; τ_*d*_ represents sensor delay; F is the control action; *x*_*ref*_ is the reference position; and *v*_*ref*_ is the reference speed.

Additive disturbance sources were included at the reference motion command and at the control action, denoted as *d*_ω_ and *d*_*f*_ respectively. Both disturbances were modeled as white noise. A constant delay of τ_*d*_ = 100 *ms* was included in the feedback path to account for feedback delays in the human sensorimotor system. The results were robust to decreasing the delay τ_*d*_; if τ_*d*_ was set to zero the same results were observed.

An idealized transfer function for the linearized open-loop system *G*(*s*), where *s* is the Laplace variable, was designed to represent two configurations: a gravity-neutral arm *G*_*arm*_(*s*) or upright stance *G*_*stance*_(*s*):

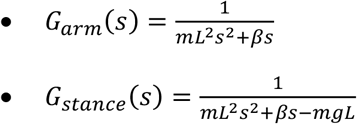

In the gravity-neutral-arm case, the system dynamics *G*_*arm*_(*s*) represents a marginally stable 2^nd^ order linear-time-invariant (LTI) system, with moment-of-inertia *mL*^2^ and damping factor β, where *m* is the total mass of the arm, and *L* is the distance between the shoulder joint and the center of mass (CoM) of the arm. The distributed moment of inertia was neglected.

In the quiet-standing case, the system dynamics *G*_*stance*_(*s*) represents an unstable 2^nd^-order LTI system i.e., an inverted pendulum, with moment of inertia *mL*^2^, damping factor β, and destabilizing gravitational stiffness *mgL*, where *m* is the total mass of a human body, *L* is the distance between the ankle joint and center of mass (CoM) of the whole body, and *g* is the gravitational acceleration (9.81 *m*/*s*^2^). Again, the distributed moment of inertia was neglected.

The closed-loop dynamics controller *C*(*s*) was designed to guarantee asymptotic stability of the closed-loop transfer function. Specifically,

- For the gravity-neutral arm, we set *C*(*s*) = *k*_*c*_ > 0;
- For upright posture, we set *C*(*s*) = *k*_*c*_ > *mgL*.

In static hand posture and quiet stance, an additional intermittent controller (IC) was introduced to bound the otherwise unbounded Brownian behavior. The chosen intermittent control performed no control action within the “do-nothing” region, while it generated a Gaussian velocity profile whenever the actual position measurement (*x*_*m*_) was outside the threshold of the “do-nothing” region. From a mathematical point of view, this was represented by the following control law:

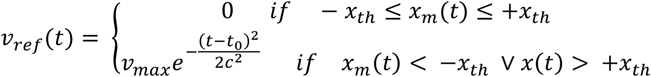

where *v*_*max*_ represents the peak velocity of the control command, *t*_0_ is the timing offset, and *c* is the standard deviation of the Gaussian profile. This intermittent control was based on the theory of dynamic motor primitives, and velocity-coded submovements specifically (Hogan and Sternad 2012). However, other control architectures that include intermittency, with a “do-nothing” region and a “do-something” region, would produce equivalent results as long as the “do-something” region included a sufficient corrective action to push the controlled system back into the “do-nothing” region.

200 simulations of this model were performed for each of the three motion tasks. The noise disturbance profiles were generated with pseudo-random white-noise sequences, initialized with different seeds to guarantee independence between the different simulations. Table 1 summarizes the values of the parameters used for the simulations. In each motor task, the position trajectory over time was sampled at a frequency of 100 Hz and then processed using the same data analysis methodology adopted for the experimental data.

**Table 1.**
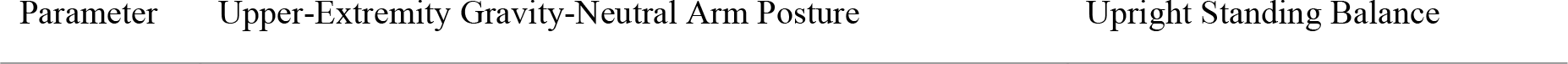
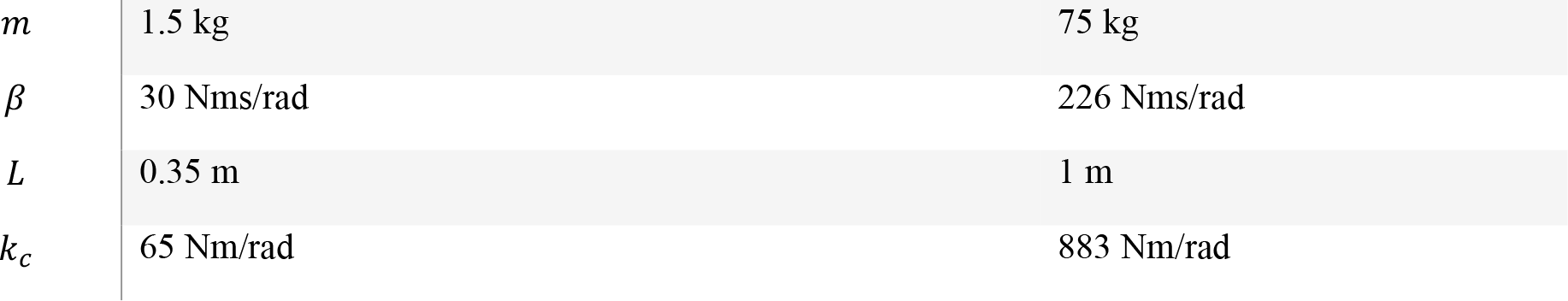
Parameters used in the simulations of the two cases considered: upper-extremity gravity-neutral arm posture and upright stance.

| Parameter | Upper-Extremity Gravity-Neutral Arm Posture | Upright Standing Balance |
| --- | --- | --- |
| $m$ | 1.5 kg | 75 kg |
| $\beta$ | 30 Nms/rad | 226 Nms/rad |
| $L$ | 0.35 m | 1 m |
| $k_c$ | 65 Nm/rad | 883 Nm/rad |

## Acknowledgments

We would like to thank the members of the Newman Laboratory for Biomechanics and Human Rehabilitation for the continuous support and valuable feedback throughout this research project.

The authors contributed to the different parts of this work in the following way (FT: Federico Tessari, JH = James Hermus, RSD = Rika Sugimoto-Dimitrova, NH = Neville Hogan):

- Conceptualization: FT, JH, RSD, NH
- Methodology: FT, JH, RSD, NH
- Investigation: FT, JH, RSD
- Visualization: FT, JH, RSD
- Funding acquisition: NH
- Project administration: NH
- Supervision: NH
- Writing – original draft: FT
- Writing – review & editing: FT, JH, RSD, NH

## Supplementary Material

### Spectral Analysis

Power spectral densities (PSDs) were computed for each data set using Welch’s method (Solomon, Jr, O M 1991). Various data pre-processing techniques were compared with the goal of reducing any numerical artifacts in the computed PSDs. Three different detrending approaches were considered: (I) no detrending, (II) subtraction of the mean, and (III) linear detrending. Furthermore, two windows were tested on the data: (I) rectangular window, and (II) Hanning window. Fig S1 shows PSDs computed using different detrending and windowing approaches for one subject performing the hand posture task. The different approaches mainly affected the three lowest-frequency data points, which were therefore removed from the reported PSDs. The window shape mainly affected the total signal power, without significantly affecting the shape of the spectrum. This can be explained by the fact that the Hanning window is used to taper the ends of the data window, effectively reducing signal amplitude. The Hanning window is typically used to obtain cleaner high frequency spectral estimates, reducing side-lobe artifacts that may be produced by other windowing methods, including the rectangular window. Since we were interested in the low-frequency spectrum, which is largely unaffected by the choice of the window, the final PSDs were computed using the rectangular window as it involved minimal processing of the data.

**Fig S1.**
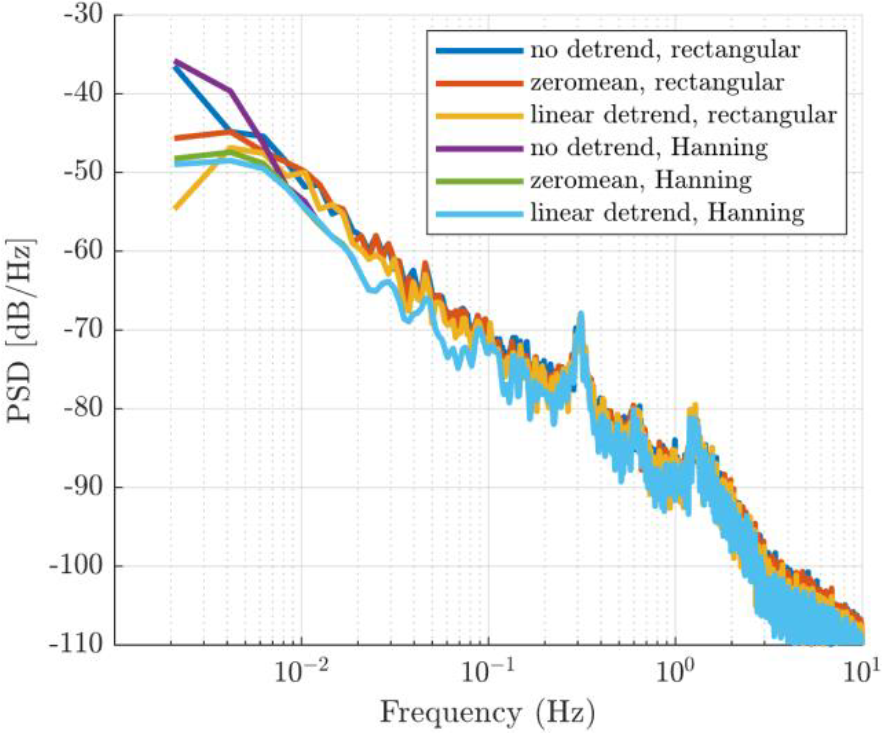
Power spectral densities for a single subject performing the hand posture experiment. They were computed using six different pre-processing techniques: no detrending with rectangular window (blue), zero-mean with rectangular window (red), linear detrend with rectangular window (yellow), no detrending with Hanning window (purple), zero-mean with Hanning window (green), linear detrend with rectangular window (light blue). The detrending process mainly affects the lowest-frequency data points, while the window shape affects total signal power.

For the crank-turning task, which involved displacements due to voluntary movement of the hand around the crank, the crank-angle data were detrended by removing the average trajectory over the 21 trials in order to focus the spectral analysis on the random fluctuations around the average trajectory.

### Alternative Models

Two alternative models were investigated to describe a Brownian process in position emerging from a closed-loop negative feedback control system. The first was based on work proposed by Peterka (Peterka 2000), where stationary noise is filtered through a low-pass filter with an extremely low cut-off frequency, a so-called ‘leaky integrator’, and injected as a disturbance into the feedback control loop. Stationary noise filtered through a ‘leaky integrator’ generates a Brownian-like behavior that is bounded, with variance growing linearly over time until the low cut-off frequency of the filter is reached. Such a model reproduces the Brownian-like behavior observed in postural tasks, such as hand posture control or quiet standing, in which the variance is bounded. However, it fails to reproduce the unbounded Brownian behavior observed in velocity-based tracking tasks.

A second alternative model describes the Brownian behavior in position as emerging from the closed-loop dynamics. In this case the closed-loop transfer function requires exactly one pole at the complex-plane origin i.e., a free integrator. The controller required to produce this behavior differs depending on the open-loop dynamics G(s) of the system being considered, which can be:

I. unstable (at least one pole with positive real part);

II. asymptotically stable (all poles with negative real parts);

III. marginally/neutrally stable (one pole at the origin).

For cases (I) and (II), the controller *C*(*s*) was designed to cancel all unstable poles, and place one pole at the origin. In case (III), the open-loop system already presents a pole at the origin, so the controller *C*(*s*) was designed such that the pole at the origin was not shifted. The following presents three different 2^nd^ order LTI systems e.g., mass-spring-damper systems, with *G*(*s*) in each of the three previously mentioned cases, and the corresponding controller *C*(*s*) required to place a single closed-loop pole at the origin:

I. If 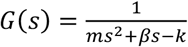, then*C* (*s*) =*k*_*c*_ = *k*

II. If 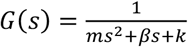, then*C* (*s*) =*k*_*c*_ = −*k*

III. If 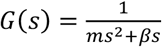, then*C* (*s*) =*k*_*c*_ = 0

The 2^nd^ alternative model, with the three variations described above to produce a free integrator, would guarantee the appearance of an unbounded Brownian behavior in position. To bound the Brownian behavior in postural tasks (hand posture and quiet standing), an intermittent controller was designed such that the control actions producing the free integrator were augmented to limit the position variance. In the postural tasks, the condition was a threshold in the position error: when the position error was within the thresholds, the closed-loop dynamics maintained a pole at the origin, while when the error exceeded the thresholds, the closed-loop dynamics exhibited an asymptotically stable behavior. Fig S2 presents a possible intermittent control action *C*(*s*) for the three considered open-loop dynamics: unstable, asymptotically stable, and marginally stable.

**Fig S2.**
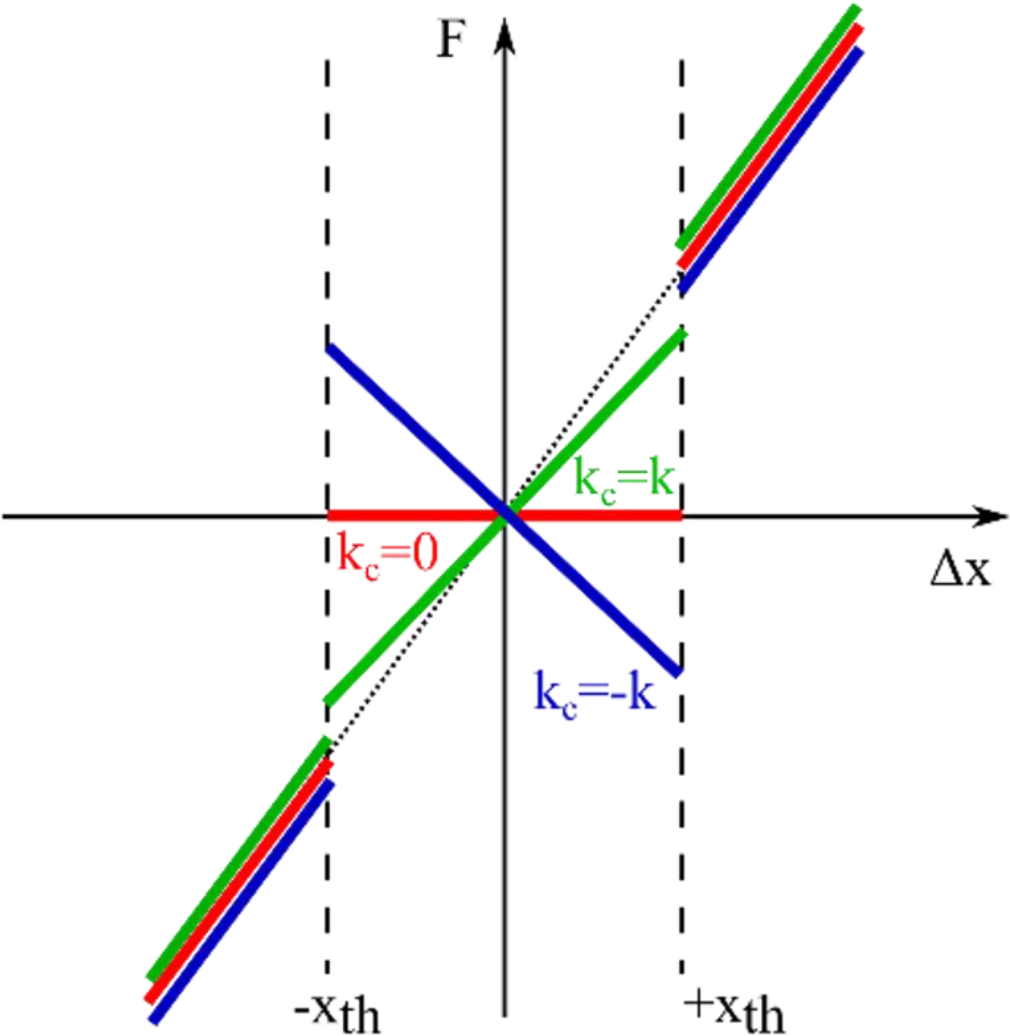
Intermittent control action based on position thresholds ±*x*_*th*_. (black dashed lines). It was designed to guarantee a free integrator in the closed-loop dynamics when the error was within the thresholds for three different open-loop dynamics: unstable (green solid line), marginally stable (red line), asymptotically stable (blue). Outside the thresholds the controller guarantees an asymptotically stable behavior.

### Stabilogram Diffusion Analysis

Stabilogram diffusion analysis was introduced by Collins and De Luca to study the fluctuations observed in the trajectory of the center of pressure (CoP) position during quiet stance (Collins and de Luca 1993). It is a time-series analysis tool borrowed from statistical mechanics that involves computing the mean-squared displacement (MSD) of a signal for different time intervals to analyze the nature of the fluctuations present in the time series.

Consider a signal sequence *x*(*k*) with *k* = 1, …, *N*, and let Δ*t* denote the span interval between two points of the *x*(*k*) data sequence. The mean-squared displacement can be computed as:

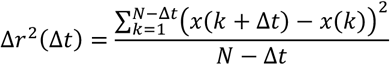

The mean-squared displacement can then be plotted with respect to different span intervals Δ*t* to produce a diffusion plot. The evolution of Δ*r*^2^ provides information on the fractality of the analyzed time series. In fact, it is known that a fractional random walk process is characterized by the following relationship between the mean-squared displacement and time:

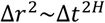

where the scaling exponent *H* denotes the Hurst coefficient, and can be obtained from a log-log plot of Δ*r*^2^ with respect to Δ*t*. When the slope 2H is exactly equal to 1, corresponding to *H* = 0.5, the behavior follows a classical (purely) Brownian process, with variance that grows linearly with time. Instead, if *H* ≠ 0.5, the process is classified as a fractal or “fractional Brownian” process, which can be persistent (*H* > 0.5) or anti-persistent (*H* < 0.5) (Collins and de Luca 1993).

Collins and De Luca observed that their diffusion plots exhibited two distinct regions with different slopes: what they referred to as a “short-term” region and a “long-term” region. The slopes measured in the log-log diffusion plots suggested the presence of “fractional Brownian” processes in both regimes, with persistent behavior in the short term, and anti-persistent behavior in the long term. Furthermore, the change in slope, from a persistent to an anti-persistent regime, was interpreted as a sign of intermittent control. However, it is important to observe that the system considered here is an inertial system, and that may account for higher-order trends manifest in the diffusion plot, including a change in the observed slope. By means of numerical simulations in MATLAB, we observed – Fig S3 – that it is possible to reproduce a diffusion plot like that of Collins and De Luca, with two regions with different slopes.

**Fig S3.**
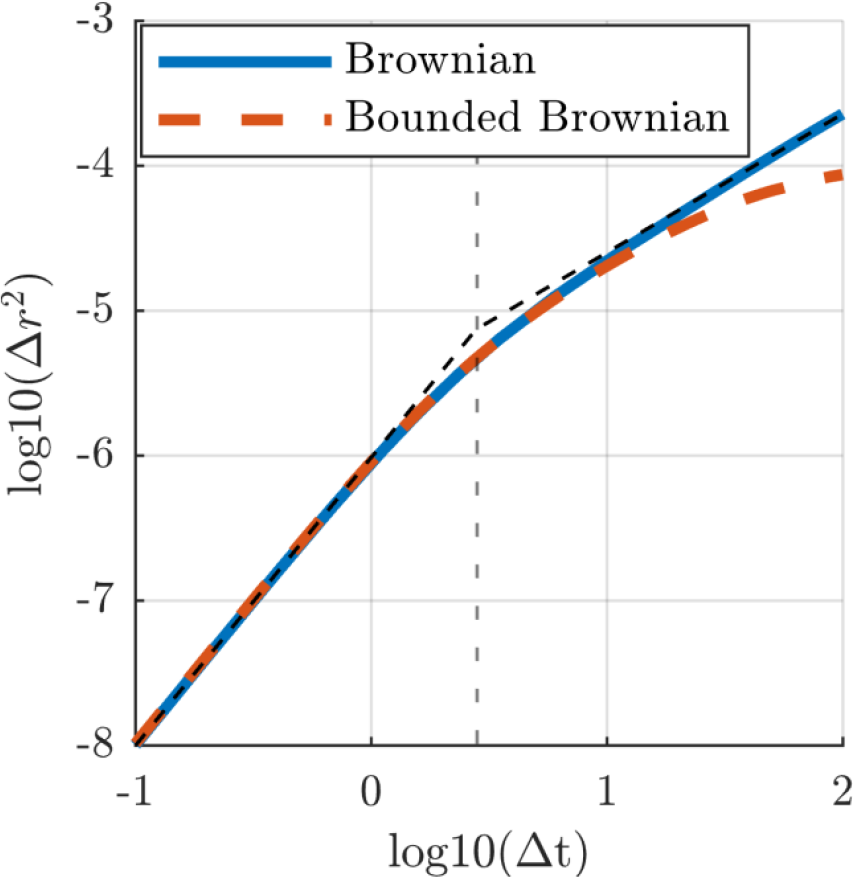
Diffusion plots of a Brownian process (blue line) and of a bounded Brownian process (dashed red line). Both were filtered through a second order mechanical system. The vertical dashed grey line indicates the critical point that divides the short-term and long-term regions with different slopes. The black dashed lines represent the best-fit lines to the Brownian process before and after the critical point.

We considered two different inputs: (i) a purely Brownian input and (ii) a bounded Brownian input, obtained by bounding the Brownian process. We then injected these noise sources into a 2^nd^ order dynamical system comprising a mass, spring and damper. The results – presented in Fig S3 – show the average of 100 simulations performed considering a pseudo-random white noise source with randomized initial seed integrated to obtain a purely Brownian time series that was filtered through a 2^nd^ order mechanical system with the following properties: 75 kg mass, 245 Ns/m damping factor and 200 N/m stiffness. When we consider a purely Brownian process input (blue line of Fig S3), the slope of the long-term region is exactly 1, corresponding to *H* = 0.5. On the other hand, when a bounded Brownian input (dashed red line of Fig S3) was injected in the system, the slope of the long-term region decreased such that *H* < 0.5. The slope of short-term region was 2 (*H* = 1) with both inputs (purely Brownian and bounded Brownian), and likely arose from inertial dynamics, which is expected to exhibit “persistent” behavior. These results also agree with the spectral analysis of the experimental data: the Brownian process dominates at lower frequencies (long time scales), with flatter slopes observed when the process is bounded.

### Confidence Intervals

A linear regression model was used to fit the low-frequency region of the PSDs of the analyzed motor tasks. The linear regression was performed in the log-log plane, on the Bode magnitude plot of the PSDs. For each subject a best-fit slope *m*_*sl*_ [dB/dec] and a best-fit intercept *q*_*sl*_ [dB] was identified such that:

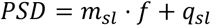

Additionally, for each regression line, the coefficient of determination *R*^2^ was computed to assess how well a linear model fit the low-frequency data.

The confidence intervals (CI) on the identified slope *m*_*sl*_ of the PSD were computed using the following equation:

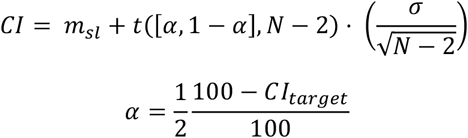

where *t*([α, 1 − α], *N* − 2) represents the t-score cumulative distributed function values for a confidence range of α. *N* represents the number of samples, while σ is the standard deviation of the slope [dB/dec]. We set the confidence interval target at *CI*_*target*_ = 95%.

The standard deviation of the slope σ was computed as:

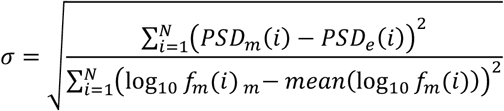

where *PSD*_*m*_ is the measured power spectral density, *PSD*_*e*_ is the estimated power spectral density obtained from the linear regression model, and *f*_*m*_ is the measured frequency.

The data presented in the main text showed the performance of only one subject for each analyzed condition. Here we report the 95% confidence intervals (CIs) for the identified slope *m*_*sl*_ for the Bode-magnitude PSD of each subject for each motor tasks: crank turning (Fig S4), hand posture (Fig S5), and quiet standing (Fig S6). Moreover, Table S1, Table S2 and Table S3 report respectively the *R*^2^ coefficients for the PSD linear fitting of crank turning, hand posture and quiet standing.

**Fig S4.**
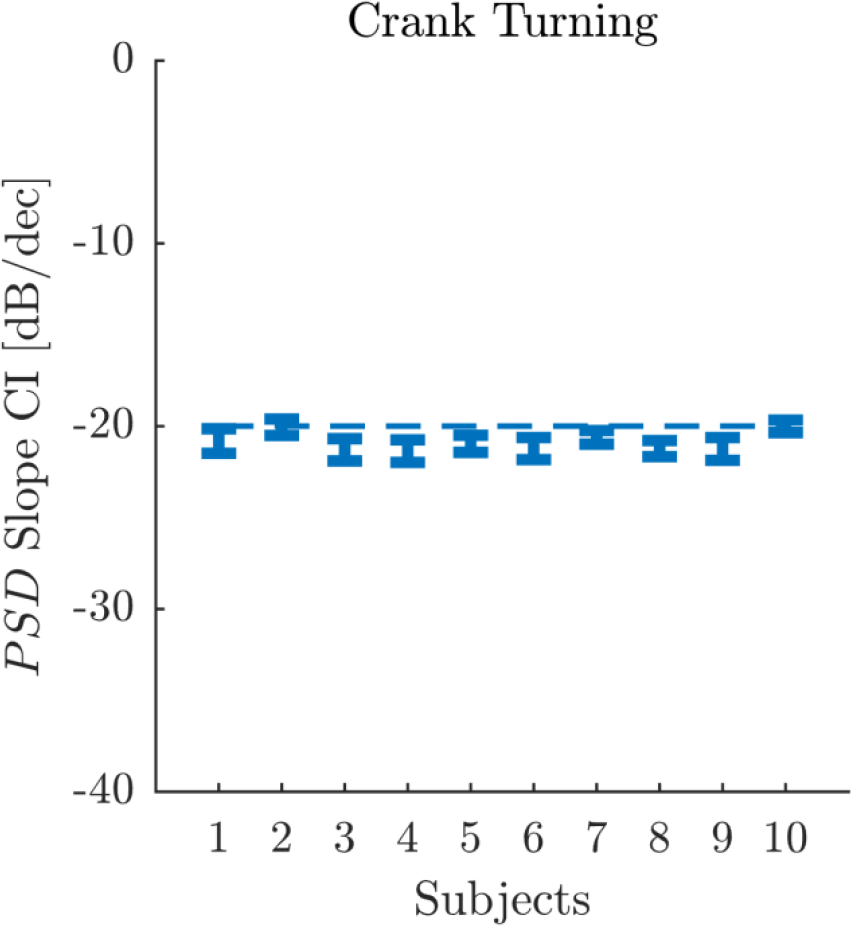
Confidence Intervals (CI) of the best-fit slope for the Bode-magnitude PSD of the crank angular position for each tested subject during the crank-turning experiment.

**Fig S5.**
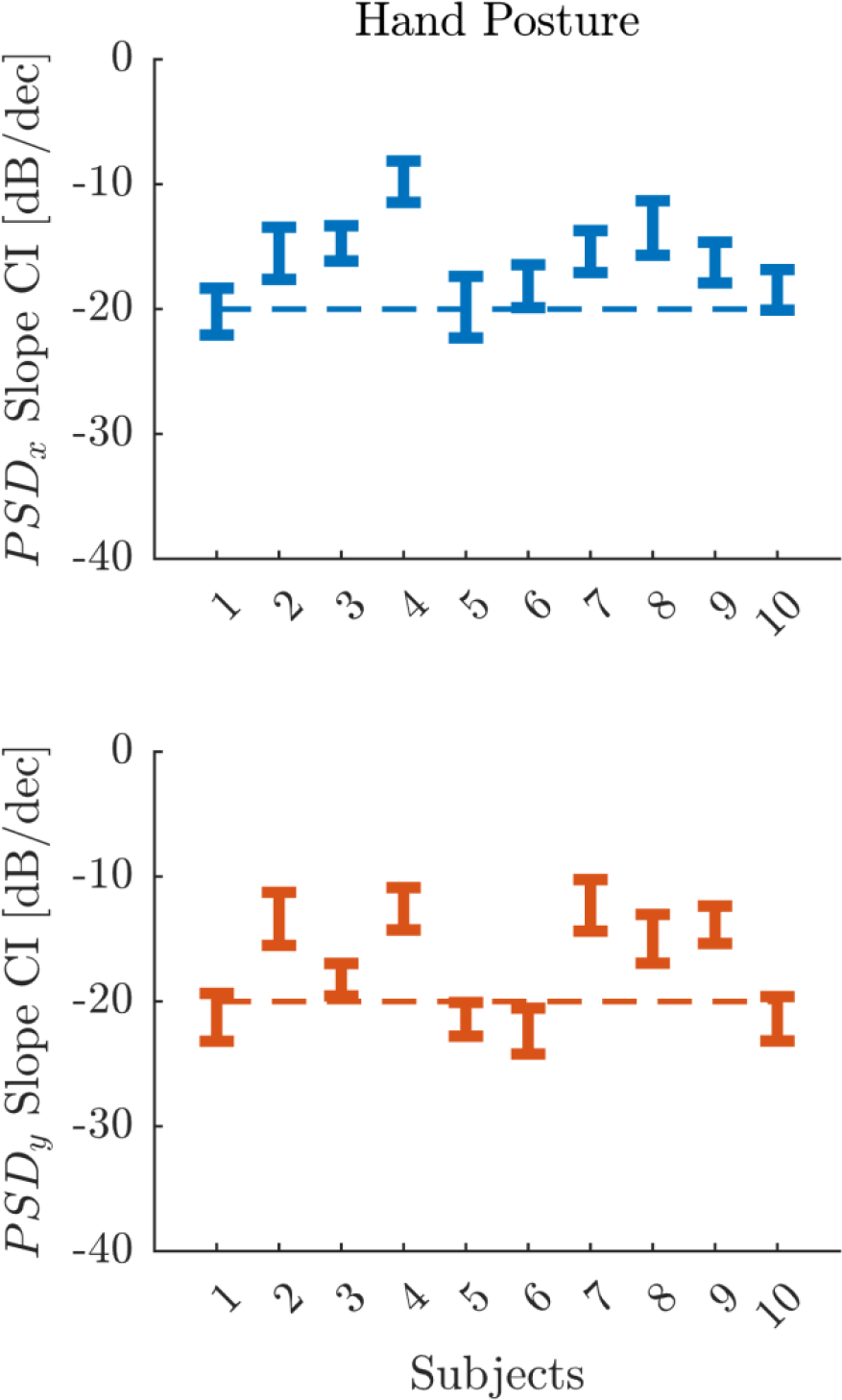
Confidence Intervals (CI) of the best-fit slope for the Bode-magnitude PSD of the hand Cartesian position for each tested subject during the hand-posture experiment. The top chart shows the ‘x’ component of the hand trajectory, while bottom chart shows the ‘y’ component of the hand trajectory.

**Fig S6.**
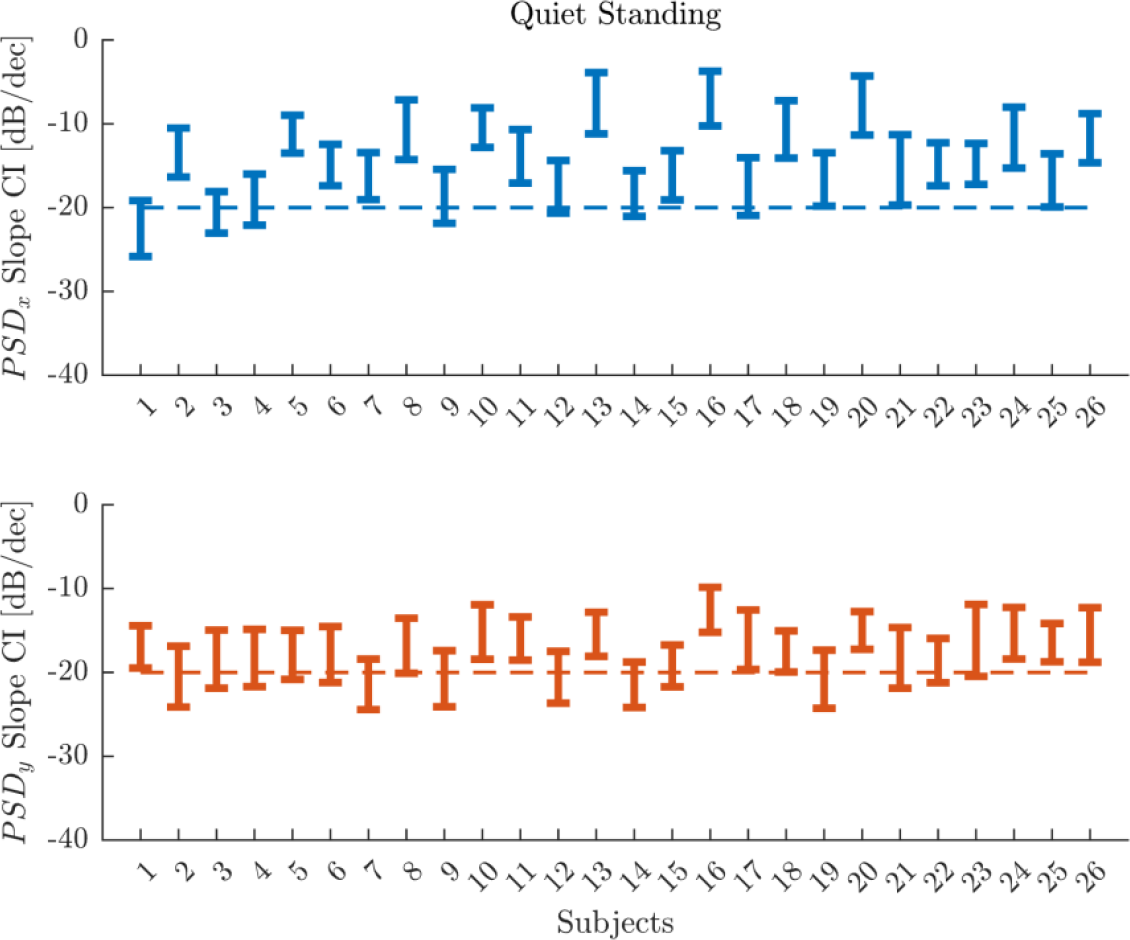
Confidence Intervals (CI) of the best-fit slope for the Bode-magnitude PSD of the center of mass (CoM) trajectory for each tested subject during the quiet standing experiment. The top chart shows the ‘x’-component of the CoM position, while bottom chart shows the ‘y’-component of the CoM position.

**Table S1.**
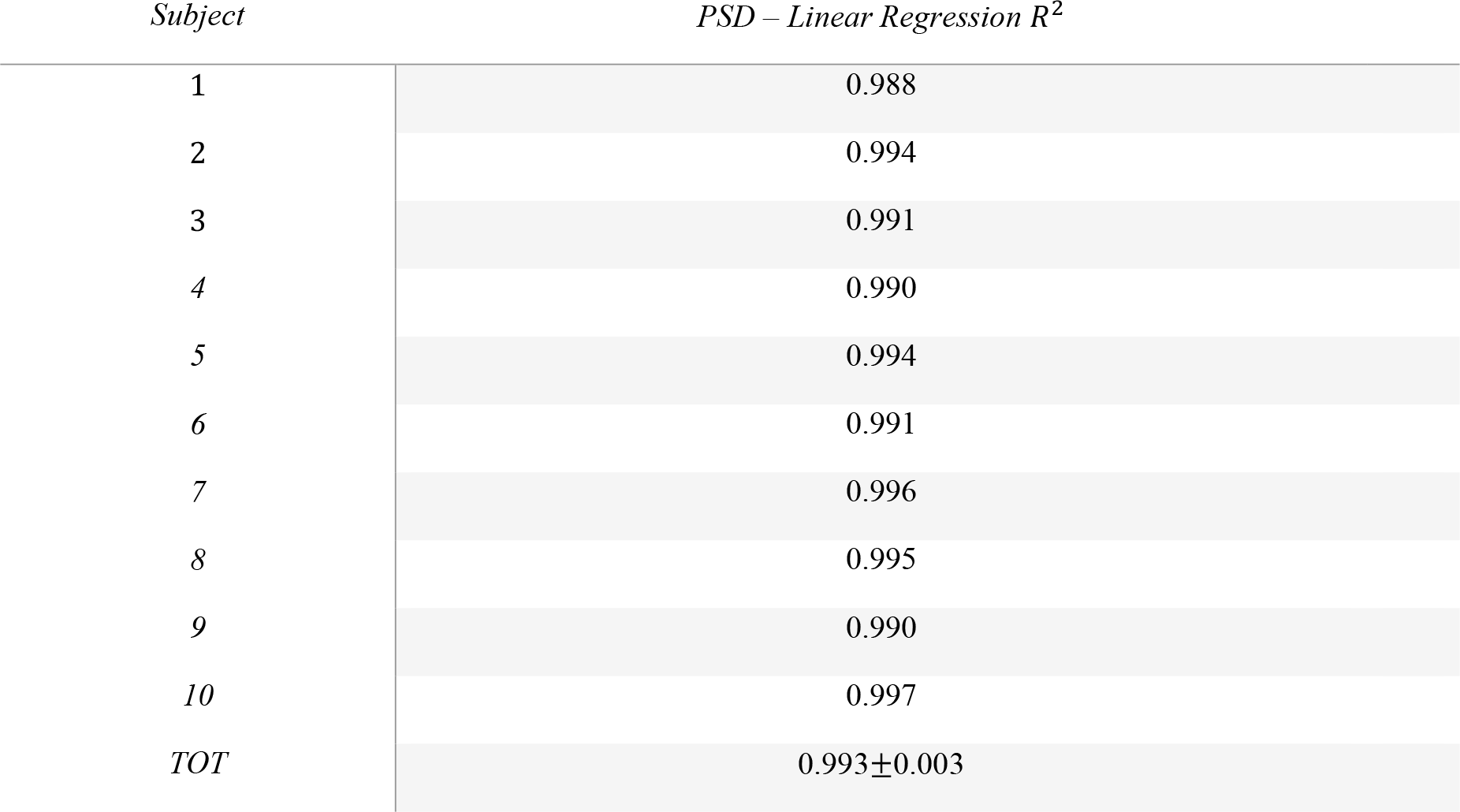
Linear regression *R*^2^ coefficients of the low-frequency PSD fit during crank turning in different subjects.

| *Subject* | *PSD – Linear Regression $R^2$* |
| --- | --- |
| 1 | 0.988 |
| 2 | 0.994 |
| 3 | 0.991 |
| 4 | 0.990 |
| 5 | 0.994 |
| 6 | 0.991 |
| 7 | 0.996 |
| 8 | 0.995 |
| 9 | 0.990 |
| 10 | 0.997 |
| *TOT* | 0.993±0.003 |

**Table S2.** Linear regression *R*^2^ coefficients of the low-frequency PSD fit during hand posture in the different subjects for both ‘x’ and ‘y’ directions.

| *Subject* | *Direction* | *PSD – Linear Regression $R^2$* |
| --- | --- | --- |
| 1 | *x* | 0.92 |
|  | *y* | 0.92 |
| 2 | *x* | 0.80 |
|  | *y* | 0.74 |
| 3 | *x* | 0.92 |
|  | *y* | 0.95 |
| 4 | *x* | 0.65 |
|  | *y* | 0.75 |
| 5 | *x* | 0.87 |
|  | *y* | 0.96 |
| 6 | *x* | 0.92 |
|  | *y* | 0.94 |
| 7 | *x* | 0.86 |
|  | *y* | 0.72 |
| 8 | *x* | 0.79 |
|  | *y* | 0.85 |
| 9 | *x* | 0.91 |
|  | *y* | 0.89 |
| 10 | *x* | 0.93 |
|  | *y* | 0.94 |
| *TOT* |  | 0.86±0.09 |

**Table S3.** Linear regression *R*^2^ coefficients of the low-frequency PSD fit during quiet standing in the different subjects for both ‘x’ and ‘y’ directions.

| *Subject* | *Direction* | *PSD – Linear*<br>*Regression $R^2$* | *Subject* | *Direction* | *PSD – Linear*<br>*Regression $R^2$* |
| --- | --- | --- | --- | --- | --- |
| 1 | *x* | 0.81 | 14 | *x* | 0.81 |
|  | *y* | 0.81 |  | *y* | 0.86 |
| 2 | *x* | 0.67 | 15 | *x* | 0.74 |
|  | *y* | 0.75 |  | *y* | 0.85 |
| 3 | *x* | 0.87 | 16 | *x* | 0.30 |
|  | *y* | 0.73 |  | *y* | 0.67 |
| 4 | *x* | 0.79 | 17 | *x* | 0.71 |
|  | *y* | 0.73 |  | *y* | 0.66 |
| 5 | *x* | 0.70 | 18 | *x* | 0.48 |
|  | *y* | 0.78 |  | *y* | 0.83 |
| 6 | *x* | 0.78 | 19 | *x* | 0.72 |
|  | *y* | 0.73 |  | *y* | 0.77 |
| 7 | *x* | 0.76 | 20 | *x* | 0.32 |
|  | *y* | 0.83 |  | *y* | 0.81 |
| 8 | *x* | 0.46 | 21 | *x* | 0.56 |
|  | *y* | 0.71 |  | *y* | 0.71 |
| 9 | *x* | 0.76 | 22 | *x* | 0.76 |
|  | *y* | 0.78 |  | *y* | 0.82 |
| 10 | *x* | 0.65 | 23 | *x* | 0.78 |
|  | *y* | 0.67 |  | *y* | 0.57 |
| 11 | *x* | 0.64 | 24 | *x* | 0.49 |
|  | *y* | 0.78 |  | *y* | 0.70 |
| 12 | *x* | 0.75 | 25 | *x* | 0.72 |
|  | *y* | 0.81 |  | *y* | 0.83 |
| 13 | $x$ | 0.29 | 26 | $x$ | 0.60 |
| | $y$ | 0.77 | | $y$ | 0.68 |
|  | TOT |  |  |  | 0.70±0.14 |

### Linear Regressions of Variance

For each subject in the crank-turning and hand-posture data, the variance trend with respect to time was computed across trials – see the Methods section for details of the computation. For each motor task and for each subject, a linear regression model was fit to the variance. In the crank-turning task, the linear model was fit over the entire trial duration. In the hand-posture task, two best-fit lines were fit: (i) one for the initial linearly growing variance, until the breakpoint was reached (*t*_*bp*_), and (ii) a second one for the entire trial duration. The breakpoint time (*t*_*bp*_) was identified as the time presenting the maximum linear fitting performance. This was found by computing the linear regression for increasing time windows and identifying the time window at which *R*^2^ reached a maximum value, after which it declined. The search was performed in increments of Δ*t* = 1 *s*, spanning from a minimum window of 1 *s* to a maximum window of 200 *s*. Tables S4 and S5 report the *R*^2^ coefficients for each subject respectively in the crank-turning and hand-posture tasks. Table S5 also reports the breakpoint time for each subject.

**Table S4.** Linear regression *R*^2^ coefficients for fits to the variance over time during crank turning in different subjects.

| *Subject* | *Variance – Linear Regression $R^2$* |
| --- | --- |
| 1 | 0.96 |
| 2 | 0.96 |
| 3 | 0.96 |
| 4 | 0.97 |
| 5 | 0.98 |
| 6 | 0.96 |
| 7 | 0.96 |
| 8 | 0.99 |
| 9 | 0.91 |
| 10 | 0.98 |
| *TOT* | 0.96±0.02 |

**Table S5.**
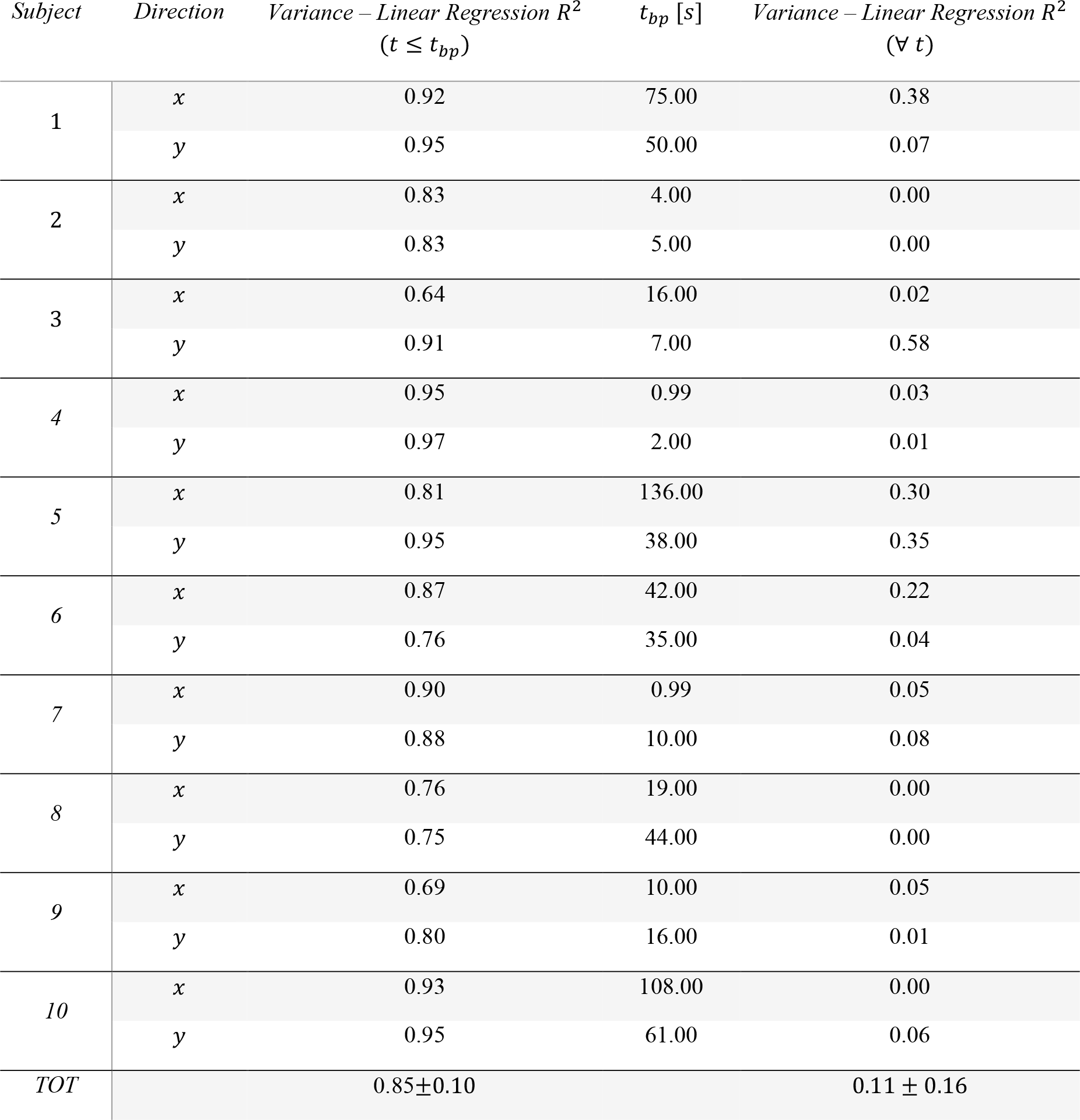
Linear regression *R*^2^ coefficients for fits to the variance over time during the hand-posture task in the different subjects for both ‘x’ and ‘y’ directions. The table reports the *R*^2^ for both the initial part with linearly growing variance until the breakpoint (*t*_*bp*_) and for the variance over the entire trial duration.

| *Subject* | *Direction* | *Variance – Linear Regression $R^2$*<br>*($t \leq t_{bp}$)* | *$t_{bp}$ [s]* | *Variance – Linear Regression $R^2$*<br>*($\forall t$)* |
| --- | --- | --- | --- | --- |
| 1 | *x* | 0.92 | 75.00 | 0.38 |
|  | *y* | 0.95 | 50.00 | 0.07 |
| 2 | *x* | 0.83 | 4.00 | 0.00 |
|  | *y* | 0.83 | 5.00 | 0.00 |
| 3 | *x* | 0.64 | 16.00 | 0.02 |
|  | *y* | 0.91 | 7.00 | 0.58 |
| 4 | *x* | 0.95 | 0.99 | 0.03 |
|  | *y* | 0.97 | 2.00 | 0.01 |
| 5 | *x* | 0.81 | 136.00 | 0.30 |
|  | *y* | 0.95 | 38.00 | 0.35 |
| 6 | *x* | 0.87 | 42.00 | 0.22 |
|  | *y* | 0.76 | 35.00 | 0.04 |
| 7 | *x* | 0.90 | 0.99 | 0.05 |
|  | *y* | 0.88 | 10.00 | 0.08 |
| 8 | *x* | 0.76 | 19.00 | 0.00 |
|  | *y* | 0.75 | 44.00 | 0.00 |
| 9 | *x* | 0.69 | 10.00 | 0.05 |
|  | *y* | 0.80 | 16.00 | 0.01 |
| 10 | *x* | 0.93 | 108.00 | 0.00 |
|  | *y* | 0.95 | 61.00 | 0.06 |
| *TOT* |  | 0.85±0.10 |  | 0.11 ± 0.16 |

## Data Availability

All data are available in the main text or the supplementary information. The MATLAB code and the experimental data are available at the following link: https://github.com/jameshermus/brownianProcess/tree/main

